# Sex-specific recombination landscape in a species with holocentric chromosomes

**DOI:** 10.1101/2024.04.15.589577

**Authors:** Sebastian Chmielewski, Mateusz Konczal, Jonathan M. Parrett, Stephane Rombauts, Katarzyna Dudek, Jacek Radwan, Wiesław Babik

## Abstract

The rate and chromosomal positioning of meiotic recombination significantly affects the distribution of the genetic diversity in eukaryotic genomes. Many studies have revealed sex-specific recombination patterns, with male recombination typically biased toward chromosome ends, while female recombination is more evenly distributed along chromosomes, or concentrated in pericentromeric region. It has been proposed that such patterns in females may counteract meiotic drive caused by selfish genetic elements near centromeres and should not occur in species devoid of clearly defined centromeres, but evidence for this expectation is scarce. Here, we constructed a sex-specific genetic map of a species with holocentric chromosomes, the bulb mite (*Rhizoglyphus robini*), a model organism for sexual selection studies with heritable alternative male reproductive phenotypes. We found a similar recombination landscape in both males and females, with a consistent pattern of increased rates towards both chromosome ends, and a higher recombination rate in females than in males. Recombination rate was positively correlated with repeat density and negatively with gene density. Our results are consistent with the meiotic drive hypothesis and suggest that the evolution of recombination patterns is closely linked to chromosome features.

## Introduction

Intragenomic heterogeneity of recombination rate is a major determinant of genetic variance through its effect on the efficacy of both positive and purifying selection (Coop & Przeworski, 2007; Ellegren & Galtier, 2016; Webster & Hurst, 2012). Therefore, studies aiming to uncover processes shaping genetic diversity, a key parameter determining the response to selection, are likely to be incomplete without considering the genome-wide recombination landscape. Local recombination rates are shaped by a variety of factors and are often correlated with genomic features such as gene and repeat element densities, GC content, and proximity to chromosome ends (reviewed in Peñalba & Wolf, 2020; Ritz et al., 2017; Stapley et al., 2017). Chromosomal rearrangements, including inversions (Berdan et al., 2021; Wellenreuther & Bernatchez, 2018), as well as fusions and fissions (Näsvall et al., 2023), may also affect recombination rates. In addition, pronounced sex differences in recombination rates have been reported across a wide range of eukaryotic taxa (Lenormand & Dutheil, 2005; Stapley et al., 2017) with important consequences for the evolutionary dynamics of sex chromosomes (Charlesworth, 2017), the genetic architecture of complex sexually selected traits (Kasimatis et al., 2021), and the resolution of sexual conflict (Connallon & Clark, 2010).

Recent meta-analyses across a broad range of taxa have consistently demonstrated sex differences in both the overall recombination rate, referred to as *heterochiasmy*, and the spatial distribution of crossover events along chromosomes, known as *recombination landscape* (Cooney et al., 2021; Lenormand & Dutheil, 2005; Sardell & Kirkpatrick, 2020, Tan et al. 2024). In many species, female genetic maps are longer than male maps, reflecting a higher average recombination rate in females. Multiple hypotheses have been proposed to explain the evolution and direction of heterochiasmy (reviewed in Sardell & Kirkpatrick, 2020). Adaptive hypotheses suggest that reduced recombination in males may have evolved to preserve favorable allelic combinations, as males are the sex often under stronger selection during haploid (Lenormand, 2003; Lenormand & Dutheil, 2005), or diploid phase (Trivers, 1988). In contrast, several proximate, physiological explanations have been proposed to account for elevated recombination rates in females. These include a protective role against errors during prolonged meiotic arrest in oocytes (Lamb et al., 2005), weaker crossover interference in females (Petkov et al., 2007), or a pleiotropic constraint linked to recombination suppression between sex chromosomes in heterogametic sex (Haldane, 1922; Huxley, 1928).

Although many hypotheses have been proposed to explain heterochiasmy, the mechanisms determining crossover positioning within each sex remain poorly understood (Cahoon et al., 2023; Sardell & Kirkpatrick, 2020). In females, crossovers are typically distributed relatively evenly along chromosomes or are biased towards pericentromeric regions. In contrast, males tend to exhibit higher recombination rates near chromosome ends (Cooney et al., 2021; Peterson & Payseur, 2021; Sardell & Kirkpatrick, 2020). Nevertheless, exceptions exist, particularly in birds, where sex differences in pericentromeric recombination are minimal or absent (Sardell & Kirkpatrick, 2020).

Crossover positioning is thought to be under strong selective control because it regulates the amount of DNA exchanged between homologous chromosomes during meiosis. When crossovers are confined to terminal regions, only the distal ends of chromosomes are reshuffled, thereby preserving existing co-adapted allelic combinations (Veller et al., 2019). However, such a pattern might reduce generation of novel and potentially advantageous genetic variation arising from more centrally located crossovers. Additionally, centrally located crossovers are associated with a higher likelihood of chromosomal mis-segregation during meiosis (Nambiar & Smith, 2016). Despite this, many taxa exhibit increased female recombination in these regions, suggesting compensatory benefits specific to oogenesis. One such benefit may involve suppression of centromere-associated meiotic drive. Unlike spermatogenesis, which results in four viable gametes are produced, oogenesis produces only one egg and three inviable polar bodies. Chromosomes with centromeres that recruit more microtubules during meiosis I are more likely to be included in the egg, providing an opportunity for selfish genetic elements to bias their transmission (Clark & Akera, 2021; Talbert, Bayes, et al., 2009, Haig, 2010).

Brandvain & Coop (2012) proposed that increased pericentromeric recombination in females may serve as an adaptive mechanism to counteract meiotic drivers. A crossover between a drive locus and the centromere disrupts their physical linkage, ensuring equal segregation probability for all four chromatids. This minimizes the transmission advantage of selfish elements, reducing the efficacy of centromere-associated drive.

Organisms with holocentric chromosomes (lacking a localized centromere) provide a valuable framework for testing the meiotic drive hypothesis and examining the centromere’s role in shaping the evolution of sex-specific recombination landscapes. According to this hypothesis, holocentric organisms are predicted to exhibit similar recombination landscapes in both sexes due to absence of a single centromere that can be exploited by selfish genetic elements (Brandvain & Coop, 2012). Despite this theoretical prediction, empirical data on recombination in holocentrics remain limited, with research primarily limited to *Caenorhabditis elegans* and Lepidoptera (moths and butterflies). To our knowledge, no study has yet examined sex-specific recombination landscapes at fine genomic resolution in any holocentric species. In *C. elegans*, comparisons between oogenesis and spermatogenesis have been restricted to a few broad genomic intervals one on a single chromosome (Cahoon et al., 2023; Lim et al., 2008; Meneely et al., 2002). Lepidoptera are not suitable for testing the meiotic drive hypothesis, as female meiosis is achiasmatic and thus lacks crossovers entirely.

Holocentric chromosomes, characterized by a diffuse centromere, have been found in approximately 350,000 animal and plant species (Král et al., 2019). Among them, acariform mites are of particular interest due to their ecological diversity and economic significance (Krantz & Walter, 2009). This group includes important agricultural pests, such as the spider mite, *Tetranychus urticae* (Grbić et al., 2011) and the bulb mite, *Rhizoglyphus robini* (Díaz et al. 2000; Jeppson et al., 1975), as well as allergenic species like the dust mites, *Dermatophagoides spp*. (Colloff, 2010). Despite their importance, genomic resources for mites remain limited. To date, no sex-specific whole-genome linkage map has been published for any mite species. Out of more than 40,000 described acariform species (Walter & Proctor, 2013), only 14 species currently have genomes assembled to chromosome-scale resolution (NCBI Genome database). The advent of next-generation sequencing technologies offers promising tools for developing of detailed genetic maps, which can be used for anchoring genome assemblies to chromosomes (Fierst, 2015) and for accurately characterizing recombination landscapes, including potential sex-specific differences (Peñalba & Wolf, 2020).

In this study, we assembled the genome of the bulb mite, *Rhizoglyphus robini*, developed sex-specific linkage maps, and characterized the genomic features associated with local variation in recombination rate. Furthermore, we used the linkage map to generate a chromosome-scale genome assembly. Bulb mites are both economically important pests (Díaz et al., 2000), but also a convenient laboratory model (Gerson et al., 1991) widely used in evolutionary and ecological studies (e.g. Łukasiewicz et al., 2020; Plesnar-Bielak et al., 2013; Smallegange et al., 2017). It is particularly valuable for studies of sexual selection, due to the presence of two distinct male morphs differing in expression of a sexually selected trait (Radwan, 2007). Fighter males possess an enlarged third pair of legs which they use as a weapon in intrasexual competition, while scrambler males, with unmodified legs, adopt a non-aggressive strategy to obtain access to females (Radwan, 2009). A recombination map anchored to a high-quality genome assembly will facilitate identification of genomic regions underlying male morph determination-a trait with substantial additive genetic variance (Parrett et al., 2023). Previous efforts to locate these regions have been hampered by the highly fragmented genome assembly available to date (Parrett et al., 2022). Finally, the sex-specific linkage map allows us to investigate heterochiasmy and sex-specific recombination landscape in a species with holocentric chromosomes, providing an opportunity to test the meiotic drive hypothesis proposed by Brandvain & Coop (2012), which has not yet received empirical support.

## Materials and methods

### De Novo Genome Assembly and Repeat Annotation

High-molecular-weight DNA was isolated from an inbred *R. robini* IW24 line that had undergone 18 generations of full-sib mating (see Parrett et al., 2023 and Łukasiewicz et al., 2020 for inbred lines description). Approximately 13 mg of mite eggs were sequentially rinsed in 70% ethanol and Milli-Q water, transferred to a pre-cooled 1.5 mL microcentrifuge tube and gently crushed in liquid nitrogen with a metal pestle. DNA was extracted using phenol-chloroform isolation method following Sambrook & Russell (2001) and precipitated onto the glass beads supplied in the Monarch® HMW DNA Extraction Kit (New England Biolabs) to minimize DNA shearing. Beads were washed three times with the kit’s wash buffer. For short read sequencing, an additional 15 mg egg sample from the same line was processed with the DNeasy® Blood & Tissue Kit (QIAGEN). Oxford Nanopore library was prepared with the Ligation Sequencing Kit v14 (SQK-LSK114) and sequenced on PromethION using R10.4 flow cells. Illumina 2 × 150 bp libraries were sequenced on a NovaSeq X Plus platform. Nanopore reads shorter than 5 kb and with a mean Phred score below 10 were discarded using *Filtlong* v. 0.2.1 (Wick, 2018). Contaminant screening performed with *BlobTools* v. 1.1.1 (Laetsch & Blaxter, 2017) against UniProt database revealed bacterial contamination (Figure S1). To remove contaminants, only reads that mapped properly to the previously published contig-level genome assembly (Parrett et al., 2022) were retained. Mapping was done with *minimap2* v. 2.24 (Li, 2018) using the *map-ont* preset and *--secondary=no* option. Filtered reads were assembled using *Flye* v. 2.9.5 (Kolmogorov et al., 2019), with an estimated genome size of 293 Mb (Parrett et al., 2022). The draft assembly underwent three rounds of polishing with *Racon* v. 1.5.0 (Vaser et al., 2017) followed by one round of Medaka v. 2.0.1 (Oxford Nanopore Technologies Ltd., 2018). Further polishing was performed with 3 rounds of *Pilon* v. 1.24 (Walker et al., 2014), using Illumina reads trimmed with *Trimmomatic* v. 0.39 (Bolger et al., 2014). Trimmed reads were mapped to the draft assembly using *bwa-mem* v. 0.6 (Li, 2013), and duplicate reads were flagged with *samtools fixmate* and *samtools markdup* v. 1.19 (Li et al., 2009). Haplotigs and redundant contigs were removed using *purge_dups* v. 1.2.5 (Guan et al., 2020). Repeat annotation was performed using *Earl Grey* v. 5.0.0 (Baril et al., 2024), which integrates multiple tools for detection and classification of different types of repetitive elements).

### Experimental Crosses

We used inbred lines established through 10 generations of full-sib mating prior to the experiment. Four mapping families were generated by crossing mites from different lines. Mites were housed at 23°C in glass vials (∼1 cm diameter) with a plaster-of-Paris base soaked in water to maintain high humidity and fed dry yeast *ad libitum*.

To establish the parental (P) generation, juvenile mites from inbred lines were isolated into separate vials to ensure that emerging females remained virgins. Upon reaching maturity, we scored the mites for sex and male morph. After three days of mating, P individuals were transferred to DNA low-binding tubes containing DNA isolation buffer (digestion solution from the MagJET gDNA Kit, Thermo Fisher Scientific) and stored at -20°C for subsequent DNA extraction. The eggs laid during this period were left to hatch in the original vials. F1 individuals were isolated before reaching maturity to prevent uncontrolled mating, then sexed at maturity and paired with full sibs. For each of the four families, the F1 pair with the highest reproductive output was selected to continue the cross (Table 1). F2 larvae were individually separated and sexed upon reaching maturity, then preserved in DNA isolation buffer. In total, 199 *R. robini* individuals were used to construct the genetic map, including 8 P-generation founders, 8 F1 individuals, and 183 F2 progeny. The number of F2 individuals per family ranged from 32 to 60 (Table 1).

**Table 1.** Overview of the experimental crosses and their reproductive output.

| Family | dam line | sire line | F1 male morph | number of F2 individuals |
| --- | --- | --- | --- | --- |
| A | scrambler-selected IN1 | fighter-selected IW9 | scrambler | 32 |
| B | scrambler-selected IN5 | fighter-selected IW22 | scrambler | 32 |
| C | fighter-selected IW15 | scrambler-selected IN7 | scrambler | 60 |
| D | scrambler-selected IN3 | fighter-selected IW17 | scrambler | 59 |

### Whole-Genome Resequencing and Calculation of Genotype Likelihoods

Genomic DNA was extracted from whole individuals using the MagJet Genomic DNA Isolation Kit (Thermo Fisher Scientific), following the kit’s standard protocol. Genomic libraries were prepared using the NEBNext Ultra FS II kit (New England Biolabs) and resequenced on an Illumina NovaSeq 6000 platform with an S4 flow cell at the SNP&SEQ Technology Platform (Uppsala, Sweden), generating 150 bp paired-end reads Sequencing was performed to achieve an average target coverage of 20× for P and F1 individuals and 10× for F2 offspring.

Adapter trimming and removal of low-quality bases were conducted using *Trimmomatic*. Cleaned reads were mapped to our de novo *R. robini* genome assembly using *bwa-mem* with default parameters. *Samtools* was used to mark duplicate reads and retain only those with a mapping quality score above 20.

To minimize genotyping errors associated with low-coverage data, we did not call genotypes. Instead, we calculated genotype likelihoods for an initial set of SNPs using *samtools mpileup* and the *mpileup2Likelihoods* script from *Lep-MAP3* v. 0.5.1 (Rastas, 2017). To validate pedigree structure, we assessed pairwise relatedness among individuals within each family using the *Identity By Descent* (IBD) module of *Lep-MAP3*, based on a random 10% subset of markers.

### Linkage Mapping

We constructed the bulb mite genetic map using *Lep-MAP3*, following its standard pipeline. Initially, the *ParentCall2* module was used to correct missing or erroneous parental genotypes, remove non-informative markers, and identify sex-linked markers. Marker filtering was then performed using the *Filtering2* module with stringent criteria: we retained only markers with a minor allele frequency above 0.05 across families, a missing genotype rate < 25% within families, and informative (heterozygous) in all 4 families. For contigs containing fewer than 5 markers, loci heterozygous in at least 3 families were included. To conservatively remove repetitive elements and loci with non-Mendelian inheritance, we excluded markers showing significant segregation distortion (*dataTol* = 0.05), and discarded 200 bp flanking both sides of annotated repeat. Two F2 individuals were removed due to insufficient sequencing coverage (<0.1×), and one additional F2 was excluded based on high genotype missingness, as determined using *PLINK* v. 1.9 (Purcell et al., 2007; see Supplementary Material).

Markers were assigned to linkage groups using the *SeparateChromosomes2* module with LOD threshold optimized across a range of 18 to 30. We selected LOD threshold of 21 which grouped markers into eight linkage groups, consistent with the haploid chromosome number in *R. robini* (Parrett et al., 2022)

A sex-averaged map was constructed using the *OrderMarkers2* module with the Morgan mapping function, using grandparental genotypes to phase offspring genotypes. Given the XX/X0 sex-determination system in bulb mites (females carry two X chromosomes and males only one), we ordered markers on the sex chromosome using only female-informative loci (informativeMask = 2). Marker ordering was repeated 10 times, selecting the iteration with the highest likelihood for the final map.

Visualization with LMPlots revealed a single ordering error on linkage group LG8, which was corrected by reordering markers 60 times. Genotyping errors tend to cluster at the ends of linkage groups, where even a single erroneous genotype can inflate map length by introducing spurious crossovers (Cartwright et al., 2007; Rastas, 2017). To address this, we manually inspected the terminal regions of each linkage group and removed any markers disproportionally increasing genetic distance. Six markers at the end of LG8 were excluded, as these belonged to different contigs and artificially inflated the group length by 14 cM.

### Genome Anchoring

We anchored the de novo assembled *R. robini* genome using *Lep-Anchor* v. 2 (Rastas, 2020). Marker interval data from the *OrderMarkers2* output of *Lep-MAP3* were used as input. *PlaceAndOrientContigs* was used to determine the order and orientation of contigs along each chromosome, iteratively maximizing the correlation between physical and genetic distances across markers. Contig blocks without map support were removed. Linkage group labels, initially ranked by marker count, were renamed as chromosome numbers based on their physical size.

### Evaluation of the Linkage Map

We evaluated the genetic map based on the physical positions of markers after anchoring. The evaluation was performed using the *OrderMarkers2* module with the Morgan mapping function, applying parameters: *UsePhysical*, *ImproveOrder* = 0, and *EvaluateOrder* = 1, using grandparental generation data to phase genotypes. To produce sex-specific maps, we generated a female map using markers informative in mothers or in both parents (*infMask* = 23), and a male map with markers informative in fathers or in both parents (*infMask* = 13). The evaluation of the sex chromosome was based only on the female-informative markers (*infMask* = 2), consistent with the XX/X0 sex-determination system. Diagnostic plots generated with *LMPlot* identified one ordering error on chromosome 1 in both male and female maps, and an additional error on chromosome 8 in the male map. These were corrected by reordering markers with the *ProximityScale* parameter set to 100.

Genetic maps were further analyzed using custom R scripts (R Core Team, 2024). These analyses included linear modeling, correlations between recombination rates and genomic features, localization of crossover events, and evaluation of chromosomal shuffling (see below). Data visualization was carried out using the *ggplot2* package in R (Wickham, 2016).

### Recombination landscape and heterochiasmy

To investigate genome-wide patterns of recombination, we examined the relationship between chromosome size and both recombination rate and genetic map length using linear models (LMs), incorporating sex as a categorical variable to assess sex-specific effects. Recombination rates were calculated in 1 Mb non-overlapping windows by dividing the genetic length of each window (in centiMorgans) by its physical length (in megabases). To enable direct comparison between male and female recombination landscapes despite differences in total map length, we standardized recombination rates by dividing them by the sex-specific whole-genome recombination rates for each window. To allow consistent comparisons across chromosomes of varying length, autosome coordinates were scaled to a range of 0 to 1 by dividing each window’s position by the total length of its chromosome. The resulting sex-specific landscapes were visualized using LOESS smoothing (span = 0.75) along each chromosome.

To quantify recombination bias toward chromosome ends, we calculated a *periphery bias* following Haenel et al. (2018): recombination rate in the outermost 10% of each autosome was divided by the average recombination rate across the whole chromosome.

We investigated whether sex differences in crossover positioning contribute to variation in the amount of DNA exchanged during meiosis by calculating the probability that a random pair of loci from the same chromosome will recombine due to a crossover event (termed *intrachromosomal shuffling*, or *r̄_inter_* by Veller et al. (2019), using the following equation:

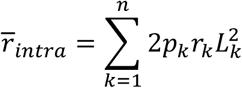

where *p_k_* and *r̄_k_* denote proportions of chromosome *k* inherited from different parental haplotypes, and *L* is the fraction of the total genome represented by chromosome *k*. Due to minor gaps in phasing, as shown in Figure S15, the sum of, *p_k_* + *p_l_* does not equal 1 (see Figure 5 in Veller et al., 2019). Finally, we estimated the probability of recombination between two loci located on different chromosomes (interchromosomal shuffling, or *r̄_inter_*) using the following equation:

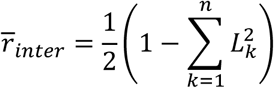

### Identification of Genomic Features Affecting Recombination Rate

To identify genomic features associated with variation in recombination rate, we examined 2 Mb non-overlapping windows across the genome and tested for correlations with sex-specific recombination rates. The genomic features analyzed included gene density, repeat density, repeat-masked GC content, and distance to the nearest chromosome end. Predicted gene annotation of the previous *R. robini* genome assembly was downloaded from *ORCAE* (Parrett et al., 2022; Sterck et al., 2012), https://bioinformatics.psb.ugent.be/gdb/Rhizoglyphus_robini), and lifted over to our new genome assembly using *liftoff* v. 1.6.3 (Shumate & Salzberg, 2021). The annotation contained an unexpectedly high number of predicted genes (∼60.000), likely reflecting pseudogenes or gene prediction artifacts. To reduce false positives, we filtered out genes with expression levels below 1 FPKM, based on transcriptomic data from Plesnar-Bielak et al. (2022), retaining a final set of 24,827 expressed genes mapped to chromosomes.

Gene and repeat densities were calculated as the proportion of each window covered by annotated features, including introns, using *bedtools coverage* v. 2.30.0 (Quinlan & Hall, 2010). GC content was estimated from the repeat-masked genome using *faCount -dinuc -strands*, and the distance from each window’s midpoint to the nearest chromosome end was calculated using a custom R script. Due to substantial collinearity among genomic features and the absence of crossovers in many windows, direct modeling of recombination rate was not feasible. Instead, we quantified associations using Spearman’s rank correlation coefficients (ρ) between each genomic feature and sex-specific recombination rate in 2 Mb windows.

## Results

### Genome Assembly

Oxford Nanopore sequencing produced 1,096,732 reads with an N50 of 19.5 kb and a mean quality score of 19.2. After filtering for a minimum mean quality of 10 and a minimum length of 5 kb 931,931 reads were retained. To reduce the risk of including contaminant sequences to draft assembly, we further selected 638,828 primary reads (i.e. aligning uniquely) to the previously published *R. robini* genome assembly (*VIB_bulbmite_20200,* Parrett et al., 2022). These reads were used for genome assembly, which after polishing and removing haplotigs resulted in obtaining the draft assembly of a total length of 293 Mb comprising 380 contigs with an N50 of 2,9 Mb and L50 of 29. Assembly completeness was assessed using BUSCO, which identified 94.80% of the 2,934 core arachnid genes detected as complete: 88.6% were detected as full-length single-copy and 6.2% as duplicated; 1.2% were classified as fragmented and 4% were missing. Repeat annotation revealed that 33% of the genome (96.19 Mb) consists of repetitive sequences. The most abundant identified categories were simple repeats (4.01%) and DNA transposons (2.28%, Table 1). A large proportion (23.76%) could not be classified into known repeat families.

### Linkage Mapping

Whole-genome resequencing of the 196 samples which met quality control criteria yielded a total of 5.6 × 10^8^ reads, of with 94% mapped to the *R. robini de novo* genome assembly (Table S1). The mean sequencing coverage was 26× in the P and F1 samples, and 11.2× in the F2 samples. After repeat masking and applying filters based on minor allele frequency, family informativeness, and segregation distortion, the number of SNP markers was reduced from 10,368,943 to 320,755, spanning 264 contigs which covered 94% of the genome. Of these, 272,619 markers (85% of the filtered SNPs) were assigned to eight linkage groups and used for linage map construction.

The bulb mite autosomal map contained 264,035 markers. The total length of the female autosomal map was 174.35 cM-approximately 55% longer than the male autosomal map, which spanned 112.4 cM (Figure 1A). We detected 340 crossovers in female meioses and 199 in male meioses. Notably, none of the linkage groups exceeded 50 cM, suggesting that a substantial proportion of crossovers were undetected (see *Discussion*). The high marker density allowed us to precisely locate recombination breakpoints, with mean inter-marker distances of 876 bp in the female map and 956 bp in the male map. The sex chromosome, which is hemizygous in males, contained 8,509 markers spanning 33.08 cM.

Overall, genome-wide recombination rate was significantly higher in female than in male map (mean female rate = 0.84 cM/Mb, mean male rate = 0.51 cM/Mb, Table 2; F = 5.96, d.f. = 1, *P* = 0.03). Linear models accounting for sex differences revealed no significant association between chromosome size and either recombination rate (F = 2.98, d.f. = 1, *P* = 0.1, Figure 1B) or genetic length (F = 1.28, d.f. = 1, *P* = 0.28, Figure 1C).

**Table 2.** Summary of the linkage map for each chromosome and sex.

| chromosome | physical length [Mb] | map length [cM] |  | number of markers |  | F:M map length ratio | recombination rate [cM/Mb] |  | periphery bias |  |
| --- | --- | --- | --- | --- | --- | --- | --- | --- | --- | --- |
|  |  | female map | male map | female map | male map |  | female map | male map | female map | male map |
| chr1 | 54.56 | 26.26 | 27.93 | 13,932 | 13,932 | 0.94 | 0.48 | 0.51 | 2.21 | 2.57 |
| chr2 | 42.07 | 26.26 | 7.26 | 61,598 | 61,605 | 3.62 | 0.62 | 0.17 | 2.16 | 2.39 |
| chr3 | 33.17 | 19.07 | 6.77 | 1,581 | 1,590 | 2.82 | 0.58 | 0.2 | 1.46 | 3.14 |
| chr4 | 30.54 | 29.17 | 16.88 | 63,320 | 63,320 | 1.73 | 0.95 | 0.55 | 2.48 | 3.64 |
| chr5 | 30.37 | 33.69 | 29.29 | 67,133 | 67,133 | 1.15 | 1.11 | 0.96 | 2.82 | 2.56 |
| chr6 (sex) | 29.7 | 33.08 | - | 8,509 | - | - | 1.11 | - | 0.49 | - |
| chr7 | 23.41 | 27.93 | 13.97 | 29,309 | 29,309 | 2 | 1.19 | 0.6 | 2.46 | 1.26 |
| chr8 | 18.2 | 11.97 | 10.3 | 27,162 | 27,162 | 1.16 | 0.66 | 0.57 | 1.06 | 0.71 |
| sum | 262.02 | 207.43 | 112.4 | 272,544 | 264,051 |  |  |  |  |  |

**Table 3.**
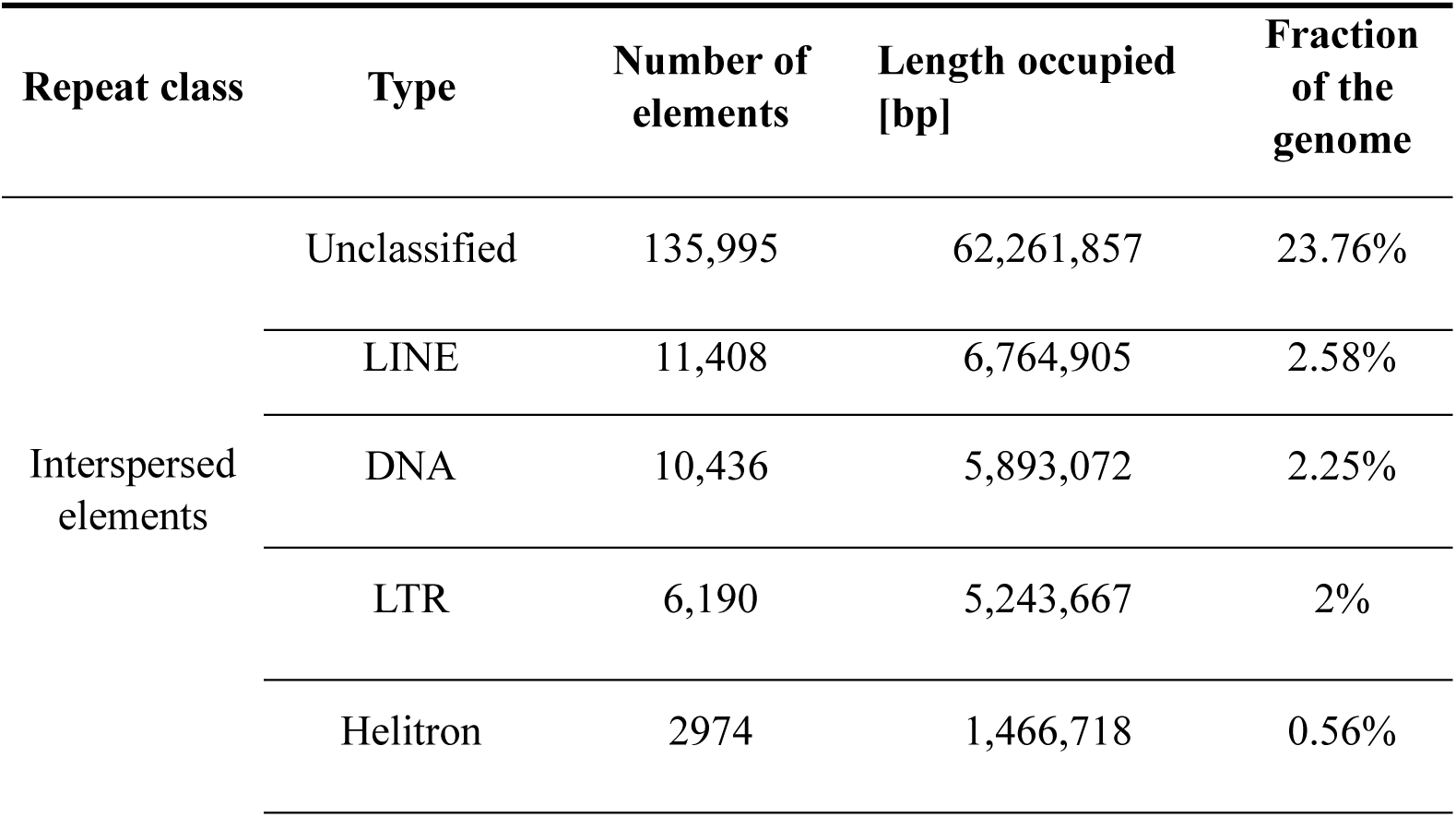
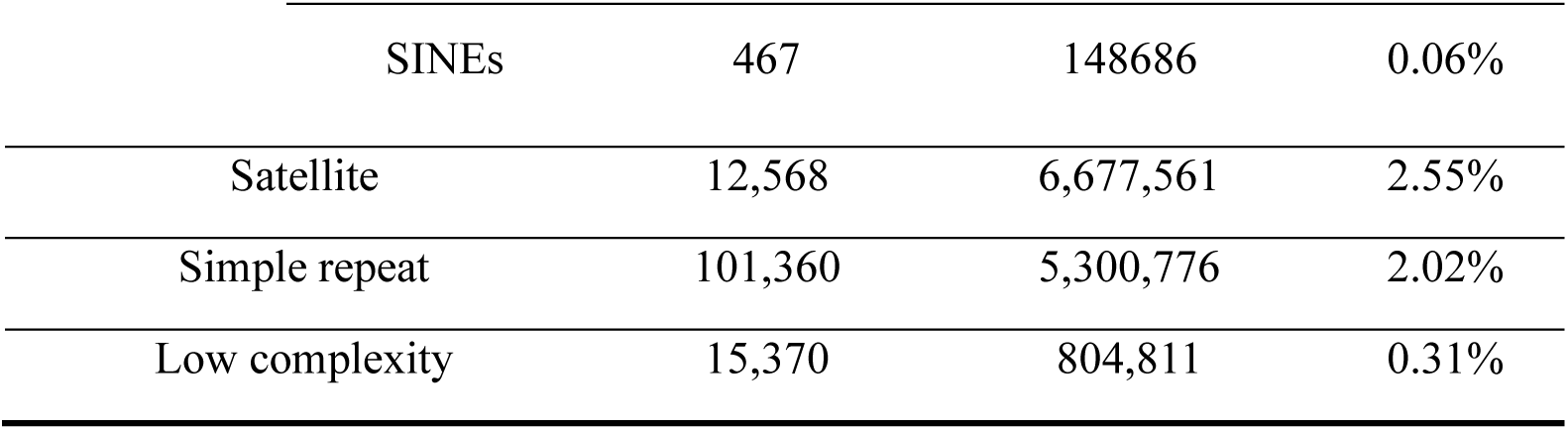
Summary of repetitive elements annotated to the major repeat classes.

### Heterochiasmy, Recombination Landscape and Allelic Shuffling

The extent of heterochiasmy varied across chromosomes (Figure 1A, Figure 1D, Table 2). Chromosome 2 exhibited the greatest sex difference in recombination rate, with female-to-male ratio of 3.62. Analysis of recombination landscapes across all autosomes revealed broadly similar patterns in males and females, characterized by elevated recombination rates toward both chromosome ends (Figure 2A). This terminally biased pattern was observed across autosomes, as indicated by the values of periphery bias exceeding one (Table 2), demonstrating an enrichment of crossovers in the outer 10% of each chromosome arm. An exception to this tendency was chromosome 8, which harbored a pronounced central crossover hotspot in males not observed on other autosomes (Figure S13). The sex chromosome also exhibited higher recombination rates in its central regions relative to chromosome termini. Periphery bias did not significantly differ between the male and female maps (LM: F = 0.85, d.f. = 1, *P* = 0.37; permutation test with 1,000 iterations: *P* = 0.36) and was not associated with chromosome length (F = 2.1, d.f. = 1, *P* = 0.17).

The estimated sex-averaged value of *r̄_intra_* in the *R. robini* genome was 0.00085. Intrachromosomal shuffling was 43% higher in females than in males (*r̄_intra females_* = 0.0010, *r̄_intra males_* = 0.0007, Figure 2B). The observed difference in*r̄_intra_* could result from differences in the overall recombination rate or from variation in crossover positioning, since crossovers occurring closer to chromosome ends reduce the effective level of genetic shuffling. However, given the no significant sex-based differences were observed in periphery bias, higher *r̄_intra_* in females is likely driven primarily by their elevated genome-wide recombination rate. We found no significant association between *r̄_intra_* and chromosome length, even when accounting for sex and including quadratic term to model potential non-linear effects (linear model: *r̄_intra_* ∼ sex + chromosome length + chromosome length²: F = 3.47, d.f. = 1, *P* = 0.089). Independent assortment of chromosomes is the prevalent mechanism responsible for allelic shuffling in *R. robini* genome, with a *r̄_inter_* value of 0.43108.

**Figure 1:**
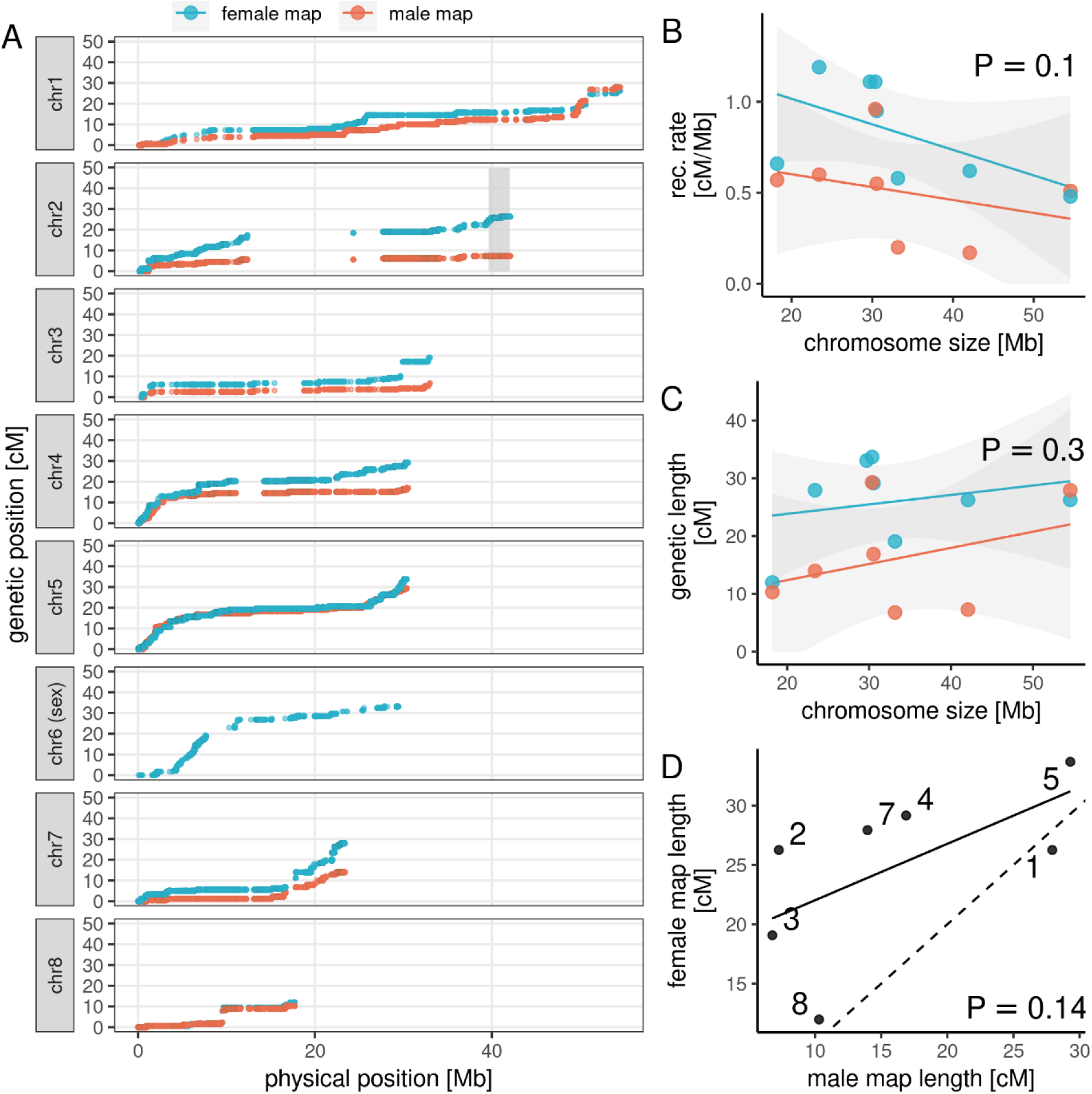
Sex-specific meiotic linkage maps and chromosome map length variation in *R. robini*. A: Marey maps displaying the genetic positions of markers in relation to their physical locations on the chromosomes. Regions with little change in genetic position, usually in chromosome centers, have low recombination rates, while steep slopes indicate high recombination rates. Grey box on chromosome 2 indicate position of contigs which diverged during artificial selection on alternative male morphs (Parrett et al., 2022), suggesting their role in the morph determination. Chromosome size was not significantly associated with either recombination rate (B) or map length (C). The solid line is the linear regression slope with 95% standard error presents as shaded areas. D: Comparison of map lengths for 7 autosomes, with female map lengths plotted against male map lengths. The dashed line indicates expected relationship under equal map lengths for both sexes.

**Figure 2:**
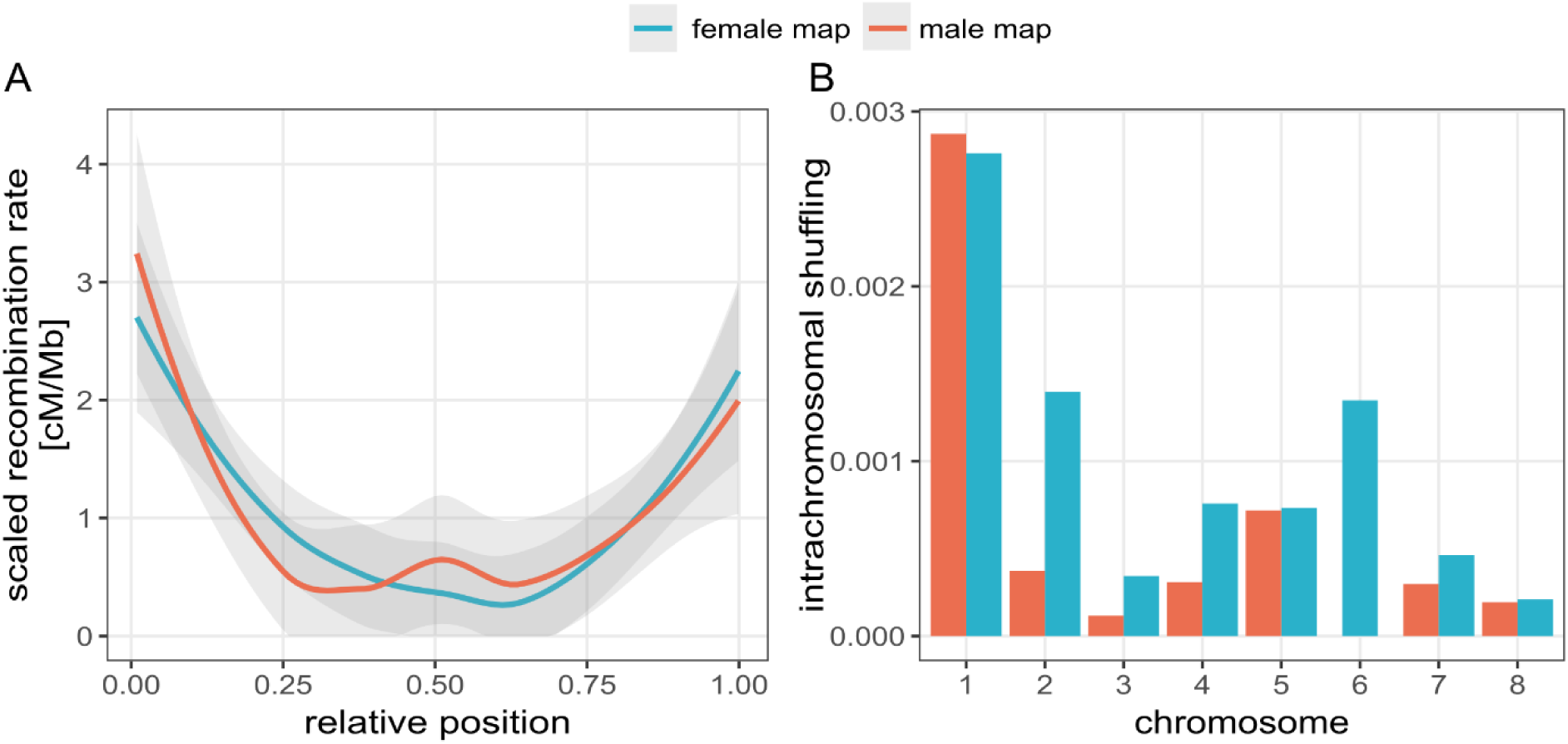
Sex-specific recombination patterns in the bulb mite genome. A: Smoothed (LOESS) line visualization of the scaled recombination rates along the relative positions of all autosomes. Only autosomes were used to provide a reliable comparison between male and female map. Scaling was performed by dividing local recombination rate by an average recombination rate of a sex-specific map. The X axis represents the relative position along the chromosomes, normalized from 0 to 1. B: Bar graph representing the average intrachromosomal shuffling value for each chromosome. Chromosome 6 is the sex chromosome.

### Anchoring the Bulb Mite genome

The linkage map was used to anchor contigs, resulting in a chromosome-level genome assembly (Figure 1). A total of 226 contigs, accounting for 89% of the genome length, were successfully anchored onto eight chromosomes. Anchoring substantially improved the genome continuity, increasing the N50 value from 2.9 Mb to 30.54 Mb. The high N50 and BUSCO scores, along with the significant integration of the genome sequence into chromosomes indicate a high-quality assembly suitable for detailed analyses of genomic features and recombination landscapes.

Chromosome sizes in *R. robini* range from 18 to 54 Mb. Gene density is highest in central regions and decreases toward chromosome ends (Figure 3). GC content follows a similar distribution, while repeat density exhibits the opposite pattern, peaking near the chromosome ends. The sex chromosome showed a significantly higher repeat content (Mann-Whitney U test; W = 1275207, *P* = 0.002) and lower gene density (Mann-Whitney U test = 1991056*, P* < 0.001) in 50 kb windows, compared to the autosomes.

**Figure 3:**
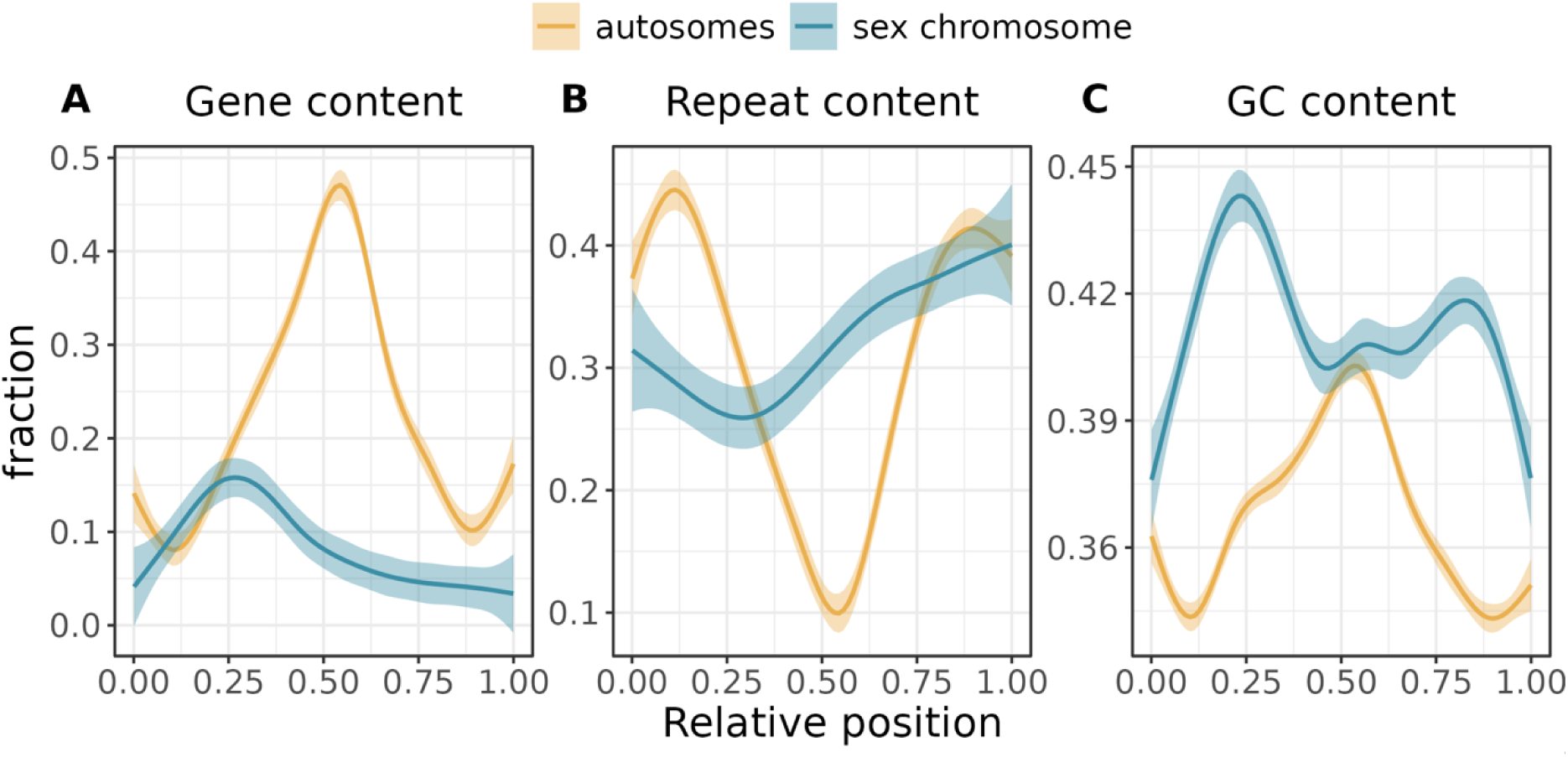
Distribution of genetic features across chromosomes in bulb mite genome. Three types of genomic features are plotted in non-overlapping 50 kb windows, which midpoints are normalized by the chromosome’s total length, to show the relative chromosomal positions, and smoothed by a LOESS line with a 95% standard error. A: The proportion of window length covered by genes (including introns), B: repetitive elements, C: GC nucleotides.

### Genomic Features Correlating with Local Recombination Rate

To identify genomic features associated with variation in recombination rate, we calculated Pearson’s correlation coefficients using 2 Mb non-overlapping windows. Recombination rate was negatively correlated with both gene density (Figure 4, Figure S12; female map: ρ = -0.36, *P* < 0.001, male map: ρ = -0.27, *P* = 0.003) and distance to the nearest chromosome end (female map: ρ = -0.53, *P* < 0.001, male map: ρ = -0.4, *P* < 0.001). In contrast repeat density was positively correlated with recombination rate (female map: ρ = 0.37, *P* < 0.001, male map: ρ = 0.3, *P* = 0.003). Recombination rates between male and female maps were strongly correlated across the genome (ρ = 0.67, *P* < 0.001), indicating highly concordant recombination landscape between the sexes. Additionally, a significant negative correlation was observed between recombination rate and repeat-masked GC content (female map: ρ = -0.39, *P* < 0.001, male map: ρ = -0.33, *P* = < 0.001).

**Figure 4:**
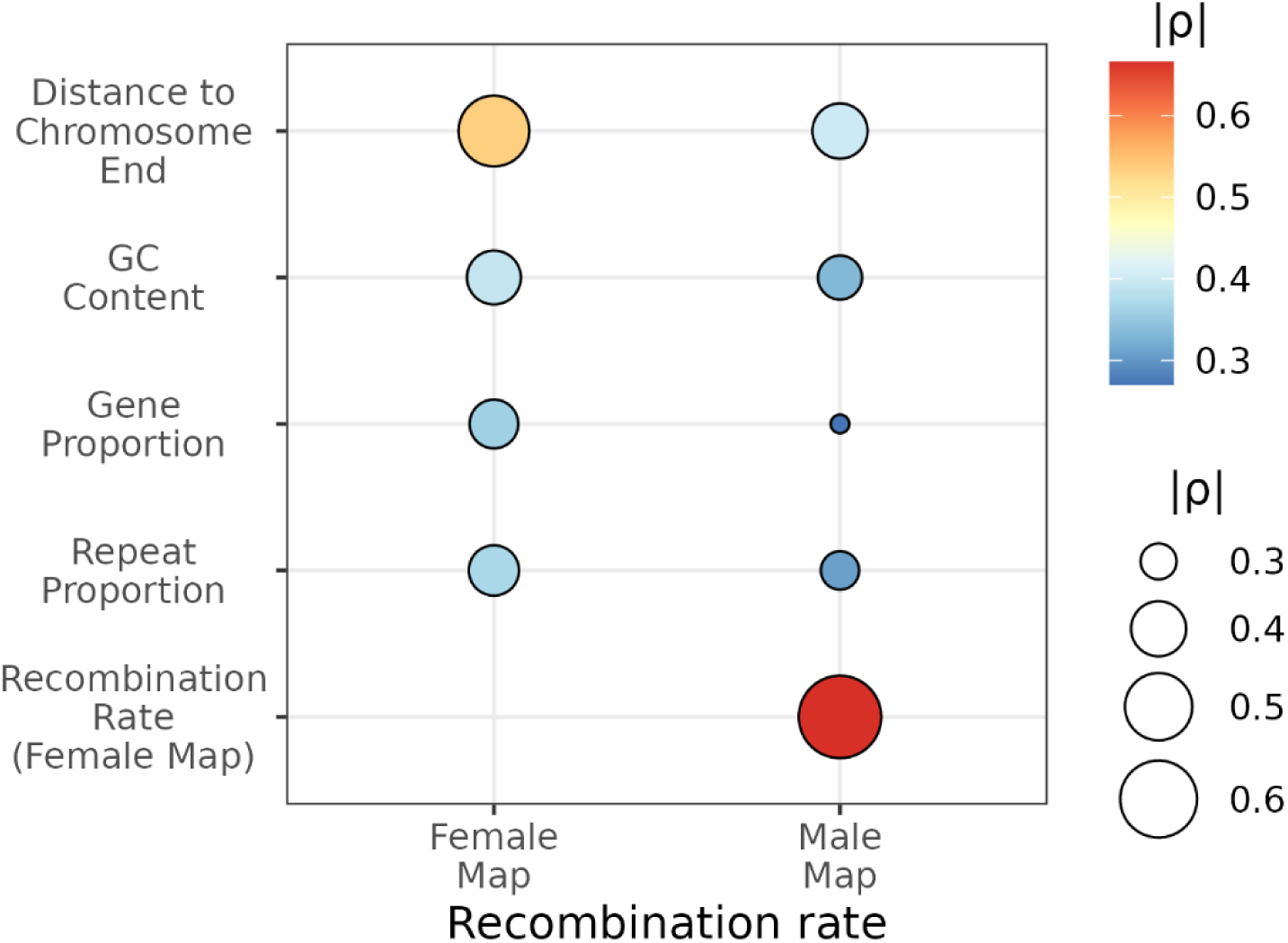
Correlation plot showing Spearman’s correlation coefficients between sex-specific recombination rates and genomic features, with the circle size indicating the strength and darkness of the color indicating the direction of the correlation. All comparisons are statistically significant (*P* < 0.05, see results).

## Discussion

The genetic map of *R. robini*, constructed using whole genomes from nearly 200 individuals, was utilized to obtain a chromosome-level genome assembly of bulb mite and evaluate the meiotic drive hypothesis regarding sex differences in the recombination landscape. To our knowledge, it is the first fine-scale sex-specific linkage map for any species with holocentric chromosomes, demonstrating that the recombination landscapes for females and males are similar. As such these results support the role of meiotic drive in the evolution of sex specific recombination landscapes.

The chromosome-scale assembly also allowed us to characterize the distribution of genomic features across *R. robini* genome. We observed a unimodal pattern of gene density, peaking centrally on chromosomes and declining toward chromosome ends and the opposite of the distribution of repetitive elements. This genome architecture, along with the elevated recombination rates at chromosome termini, resembles that of *C. elegans*, another holocentric species (The *C. elegans* Sequencing Consortium, 1998). By comparing these findings with other species, we further explore the evolutionary significance of centromeres in the evolution of sex differences in recombination landscapes.

### The Rhizoglyphus robini Genome

The genome of the bulb mite, *Rhizoglyphus robini*, spans 293 Mb, placing it among the largest genomes currently known within Acariformes. This group is generally characterized by compact genomes, with a median size of 129 Mb and a range from 32.5 to 487 Mb (Gregory & Young, 2020; Xiong et al., 2020).

Using our high-density linkage map, 89% of the genome was successfully anchored to chromosomes. Chromosome-scale assembly allowed us to identify a putative male morph determining region at the end of chromosome 2 (Figure 1A). This region contains contigs enriched for loci shown to respond to selection for alternative male morphs (Parrett et al. 2022). The co-localization of these contigs may reflect the need of coordinated expression of multiple traits required for the fighter phenotype (e.g. aggression, motor patterns and energy mobilization), which could be facilitated by the co-inheritance of alleles beneficial to fighters, but detrimental to scramblers. Such linkage could be maintained by suppressing recombination between morph-specific haplotypes, for example through chromosomal inversions, leading to formation of so-called supergenes (Mérot et al., 2020; Wellenreuther & Bernatchez, 2018). However, further studies are needed to resolve the genetic architecture underlying male morph determination in *R. robini*.

*R. robini* chromosomes exhibit a characteristic organization seen in other holocentric taxa, with gene-and GC-rich central regions and a high density of repetitive elements near the chromosome ends (Cicconardi et al., 2021; The *C. elegans* Sequencing Consortium, 1998; Zhou et al., 2023). The single X chromosome is markedly gene-poor relative to the autosomes, consistent with patterns observed in other XO sex-determination systems (Anderson et al., 2022; Y. Li et al., 2019; The *C. elegans* Sequencing Consortium, 1998). However, this reduction in gene content is not universal. In the dragonfly *Pantala flavescens*, which possesses a relatively young X chromosome, gene density on the X remains comparable to that of the autosomes (Liu et al., 2022).

Repeated sequences account for 32% of the *R. robini* genome, more than twice the proportion found in *Tyrophagus putrescentiae*, the only other acarid species for which a chromosome-scale genome assembly is currently available (Zhou et al., 2023). A large fraction (23.76%) of these repeats could not be classified into known repeat categories, indicating a potential area for future research. The elevated repeat density on the X chromosome relative to the autosomes (Figure 3C) is most likely attributable to weaker purifying selection against transposable element insertions. Under an equal sex ratio, there are four copies of each autosome but only three copies of the X chromosome in the population, because females carry two X chromosomes, whereas males carry one (Parrett et al., 2022). This reduction in effective population size amplifies Hill-Robertson interference (the selective interference among linked sites), which diminishes the removal of generally harmful TEs, allowing them to drift to higher frequencies (Kent et al., 2017).

### Sex-Specific Recombination Landscape

The genetic map of the bulb mite *R. robini* reveals pronounced sex differences in recombination rates: females exhibit rates nearly double those of males. Despite this, the overall recombination landscapes are broadly similar in both sexes, with recombination rates increasing toward chromosome ends (Figure 2A). This pattern contrasts with that observed in most monocentric taxa studied so far, where males generally show elevated recombination near telomeres, and females exhibit increased recombination near pericentromeric regions-a difference often linked to the suppression of centromere-linked meiotic drive (Brandvain & Coop, 2012; Haig, 2010; Talbert et al., 2009).

In species with monocentric chromosomes, selfish meiotic drivers can bias their own transmission into the egg by enlarging or hyperactivating the centromere to recruit more spindle microtubules during oogenesis (Clark & Akera, 2021). In response, selection in females favors crossovers between the driver and the centromere to break their physical linkage, thereby promoting increased recombination in pericentromeric regions (Haig, 2010; Talbert et al., 2009). Although such crossovers carry a higher risk of chromosome missegregation (Nambiar & Smith, 2016), they help reduce the transmission advantage of harmful drive elements. In holocentric species, such as *R. robini*, kinetochore activity is distributed along the entire chromosome, making centromere exploitation by meiotic drivers ineffective. Consequently, selection should not favor increased female recombination in pericentromeric regions, as it would not confer any benefits. Our findings of similar recombination landscapes in males and females are consistent with this prediction and support the hypothesis that sex-specific recombination patterns in monocentric species represent adaptive responses to suppress meiotic drive. However, to generalize our findings, additional sex-specific genetic maps from other holocentric taxa are necessary.

### Intrachromosomal Shuffling and Heterochiasmy

Despite broadly similar recombination landscapes in both sexes (Figure 2A), females exhibit 43% higher intrachromosomal shuffling compared to males. This difference appears to be primarily driven by a higher overall frequency of detected crossovers in females, rather than differences in crossover distribution, as supported by lack of significant sex differences in periphery bias. Intrachromosomal shuffling was not significantly associated with chromosome length, despite the fact that its calculation weights each chromosome by the square of its genomic proportion (*L^2^_k_*). This apparent paradox is resolved by strongly terminal crossover distribution: since most exchanges occur near chromosome ends, the proportions of each chromosome inherited from different parental haplotypes (*p_k_* and *r̄_k_*) remain small, reducing the influence of the chromosome size.

Heterochiasmy (the difference in recombination rates between sexes) is proposed to evolve when one sex (typically males) is under stronger selection to maintain beneficial allele combinations, leading to reduced recombination in that sex (Sardell & Kirkpatrick, 2020a; Trivers, 1988). In *R. robini,* males experience intense sexual selection through both sperm competition and physical contests, initiated by fighter males and sometimes escalating to lethality (Radwan, 1997; Radwan et al., 2000). While sexual selection might favor recombination suppression in one sex, it has also been proposed to promote increased recombination (Burt et al., 1991; Felsenstein, 1988; Maynard Smith, 1985). However, empirical studies have not consistently supported increased recombination in the sex under stronger selection (Cooney et al., 2021; Mank, 2009).

### Genome-wide and chromosomal recombination rates

With a sex-averaged recombination rate of 0.55 cM/Mb, bulb mite exhibits substantially lower recombination rate than other holocentric taxa. *Tetranychus urticae,* the closest relative of *R. robini* for which linkage map is available, shows a recombination rate of 7.63 cM/Mb (Sugimoto et al., 2020). In contrast, the more distantly related deer tick *Ixodes scapularis* was reported to have exceptionally low recombination rate of 0.006 cM/Mb (Ullmann et al. 2003). However, this estimate is based on just 127 markers across a 2.1 Gb genome, making it likely to underestimate true recombination rates, particularly if terminal markers were underrepresented, and crossovers near chromosome ends were undetected. In another holocentric group, Lepidoptera, recombination occurs only in females, yet rates still far exceed those in bulb mites, ranging from 3.6 to 7.37 cM/Mb (Näsvall et al., 2023; Torres et al., 2023).

In *R. robini,* we observed no significant relationship between chromosome size and recombination rate (Figure 1B). This contrasts with the widely reported pattern across eukaryotes, where shorter chromosomes typically exhibit higher recombination rates due to crossover assurance-the requirement that each chromosome pair undergo at least one crossover to ensure proper meiotic segregation (Brazier et al., 2025; Brazier & Glémin, 2022b; Hughes & Hawley, 2020; Jones & Franklin, 2006; Lenormand et al., 2016; Otto & Payseur, 2019; Stapley et al., 2017a; S. Wang et al., 2015). Additionally, we found no association between physical and genetic chromosome length (Figure 1C). These observations might be explained by the relatively short genetic length of linkage groups which did not exceed 50 cM-the expected genetic length when only a single crossover per bivalent is detected. Although achiasmy (absence of crossing-over) can lead to reduced total map length and has been previously reported in diplodiploid mites (Keyl, 1956), we observed crossover events on all autosomes in both sexes (Figure S14), making achiasmy an unlikely explanation for linkage groups failing to reach 50 cM. An alternative possibility is that the total map length is underestimated due to technical limitations, such as inevitable genotyping errors, which are known to affect recombination rate estimates (Lincoln & Lander, 1992; Wang et al., 2024). However, we implemented conservative strategies to minimize such biases. Instead of calling genotypes, we used *Lep-MAP3* software, which uses genotype likelihoods, allowing to account for genotyping errors, and applies error correction based on the pedigree. Additionally, we applied stringent marker filtering, removing loci with low minor allele frequency, significant segregation distortion, or overlapping with repetitive elements.

The most likely explanation for the unexpectedly short genetic map in *R. robini* lies in technical challenges associated with detecting crossovers near chromosome ends. Many species with similarly short genetic maps exhibit elevated recombination rates near telomeres, where crossover detection is particularly difficult (Backström et al., 2010; Campbell et al., 2016; reviewed in Haag et al., 2017). For example, a detailed analysis the crossover patterns in the button mushroom (*Agaricus bisporus*), based on genetic crosses and a high-quality genome assembly confirmed that the obligate crossover requirement was met (Sonnenberg et al., 2016). However, 90% of crossovers were localized within just 100 kb of chromosome ends. As a result, despite frequent recombination, the genetic map of *A. bisporus* remained exceptionally short, with an average linkage group length of only 25 cM (Sonnenberg et al., 2016). This example illustrates how terminally biased crossover distributions can lead to severe underestimation of recombination using genetic maps.

In *R. robini*, several factors likely contribute to this issue. Chromosome ends are significantly enriched in repetitive sequences, and telomeric regions were likely excluded from the assembly due to stringent marker filtering. This would have reduced marker density in telomeric regions, hampering the detection of terminal crossovers. Additionally, bulb mites possess holocentric chromosomes, which are known to exhibit terminal recombination more frequently than monocentric chromosomes (de Bigliardo et al., 2011; Heckmann et al., 2014; John, 1990; Lukhtanov et al., 2018; Nokkala et al., 2004; White, 1973; Wrensch et al., 1994). Taken together, these factors suggest that the short total map length is best explained as a technical artifact common to species in which recombination is concentrated near telomeres. Functionally, however, this underestimation is unlikely to be biologically significant, since terminal crossovers do not involve coding regions.

From an evolutionary perspective, strongly telomere-biased recombination may also be a result from reproductive transitions. Haag et al. (2017) proposed that such recombination landscape could reflect either recent transitions from asexual to sexual reproduction or introgression of asexuality-determining genes into sexual populations. In automictic species (reproducing asexually but retaining meiosis) crossovers are typically restricted to chromosome ends to minimize loss of heterozygosity, which would otherwise unmask deleterious recessive mutations and reduce fitness (Archetti, 2010; Engelstädter, 2008; Lenormand et al., 2016; Neiman & Schwander, 2011). It would be interesting to explore this hypothesis, given that the sexually reproducing *R. robini*-a member of the Acaridae family originated from oribatid mites, one of the largest known groups of asexual animals (Dabert et al., 2010; Palmer & Norton, 1992; Zhou et al., 2023). Transitions from asexuality back to sexuality have been documented in other Oribatida species (Domes et al., 2007).

### Genomic features correlating with local recombination rate

Recombination typically occurs in open chromatin and gene-rich regions, and is often positively associated with gene density, epigenetic modifications, and GC content (Arndt et al., 2005; Brazier & Glémin, 2022; Haenel et al., 2018; Kyriacou et al., 2023; Peñalba & Wolf, 2020; Stapley et al., 2017; Thuriaux, 1977). In contrast, crossover rates are usually reduced within dense heterochromatin, where transposable elements tend to accumulate, potentially disrupting gene expression and function. However, untangling the causative relationships among these covarying genomic features remains challenging.

Contrary to expectations, we demonstrated a positive correlation between recombination rate and repeat density, and a negative correlation with both gene density and distance to chromosome ends (Figure 4). These findings indicate that in *R. robini* crossovers predominantly occur in gene-poor, repeat-rich subtelomeric regions. Similarly, in holocentric organisms like *C. elegans* or some Lepidoptera, where gene density drops at chromosome ends, no discernible correlation between recombination rate and the density of coding sequence was reported (Bernstein & Rockman, 2016; Stapley et al., 2017; Talla et al., 2019; Torres et al., 2023). This suggests that the relationship between recombination and gene density may differ depending on the genome architecture.

Surprisingly, we also found negative correlation between recombination rate and GC content. This contrasts with the well-documented phenomenon of GC-biased gene conversion, which is thought to drive positive correlations between GC content and recombination rate (Bolívar et al., 2019; Duret & Galtier, 2009; Mugal et al., 2015; Webster & Hurst, 2012). In *R. robini*, this negative association may reflect the heterogeneous distribution of GC content across the genome-being highest in both gene-dense central regions and in the highly recombining subtelomeric regions (Figure 3). Among holocentric taxa, the relationship between recombination and GC content varies widely, ranging from negative (Bernstein & Rockman, 2016; Kaur & Rockman, 2014; Torres et al., 2023) to positive (Stapley et al., 2017), suggesting a complex interplay between GC-biased gene conversion and mutational biases favoring AT nucleotides (Boman et al., 2021).

The positive correlation between recombination rate and repeat abundance is likely a consequence of repeat enrichment at chromosome ends, where recombination is also elevated (Figure 3C). Typically, regions with high recombination are generally more efficient at purging deleterious TEs due to reduced Hill-Robertson interference and more effective purifying selection (Kent et al., 2017; Stapley et al., 2017). The prevalence of TEs in areas of high recombination may be attributed to their role in repairing double-strand breaks associated with recombination process (Onozawa et al., 2014), and several TE families show preferential insertion into distal parts of chromosomes (Kejnovsky et al., 2006; Pardue & DeBaryshe, 2011; Zou et al., 1996). Positive associations between recombination and specific TE families have also been reported in *C. elegans* (Duret et al., 2000), Lepidoptera (Cicconardi et al., 2021; Stapley et al., 2017b; Torres et al., 2023), and birds (Kawakami et al., 2014; Singhal et al., 2015).

## Conclusion

This study presents the first fine-scale sex-specific genetic linkage map for a holocentric organism-the bulb mite, *Rhizoglyphus robini*. Our comprehensive analyses reveal that recombination landscapes are broadly similar between males and females across most of the genome, supporting the hypothesis that elevated female recombination in monocentric species evolved as a mechanism to counteract centromere-associated meiotic drive. The absence of this pattern in *R. robini* aligns with theoretical expectations for holocentric chromosomes, where diffuse kinetochore activity precludes centromere-mediated drive. The structure of the *R. robini* genome is characterized by gene-rich central regions and repeat-rich chromosome ends, together with telomere-biased recombination, may result from their close relationship with highly asexual Oribatida. These findings offer new insights into how holocentric chromosome structure shapes recombination and open promising avenues for further research into genome evolution, sexual selection, and the genetic basis of adaptation in mites and other arthropods.

## Data availability

Raw sequence data have been deposited in NCBI Sequence Read Archive (BioProject PRJNA1088092). Code needed to repeat analyses has been deposited in GitHub (https://github.com/sebchm/genetic-map-of-the-bulb-mite) and datafiles are publicly available at Zenodo (DOI: <u>10.5281/zenodo.10959789</u>). Additionally, GFF files with annotation of genes and repeat elements have been uploaded to ORCAE (https://bioinformatics.psb.ugent.be/orcae/overview/Rhrob).

## Acknowlegements

We thank Aleksandra Łukasiewicz and Agnieszka Szubert-Kruszyńska for performing the crosses, Pasi Rastas for his generous advice on map construction and Deborah Charlesworth for her comments on the earlier version of the manuscript. Computations were partly performed at the Poznań Supercomputing and Networking Centre. This work was supported by National Science Centre grant no. 2017/27/B/NZ8/00077 awarded to J.R.

