## supplemental material for "Sex-specific recombination landscape in a species with holocentric chromosomes"

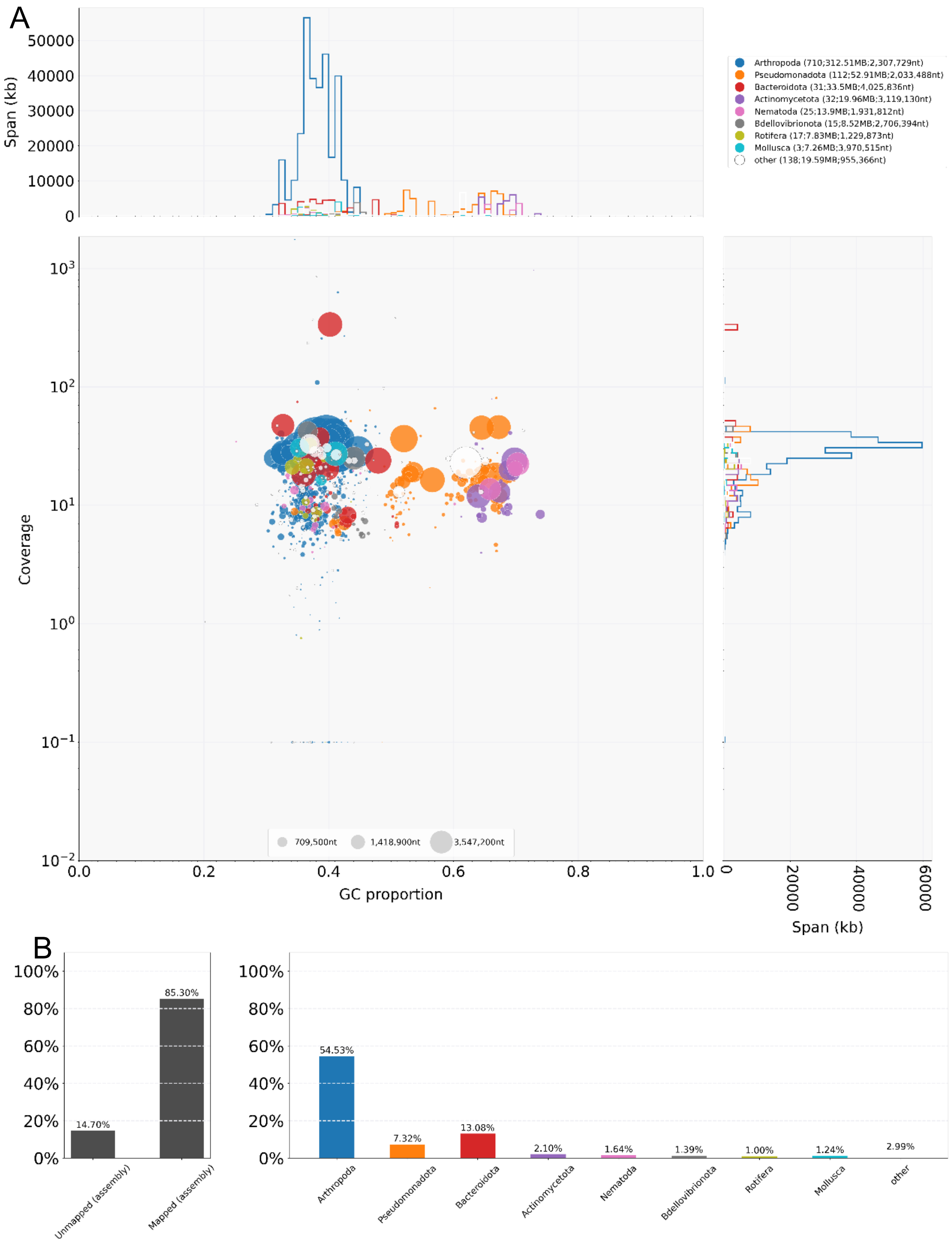


**Figure S1. Taxon-annotated GC-coverage plot of the draft *Rhizoglyphus robini* genome assembly produced by BlobTools.**

A: Each circle represents a scaffold, with diameter proportional to scaffold length and color indicating taxonomic assignment. Scaffolds are ordered by GC content (X-axis) and average sequencing coverage (Y-axis, log scale). Marginal histograms show the cumulative scaffold span for each taxonomic group across GC proportions (top) and coverage values (right). The plot highlights a clear cluster of Arthropoda-assigned sequences (blue), with additional signals from bacterial contaminants.
B: Left panel**:** Proportion of assembly mapped and unmapped contigs to known taxa in the Uniprot database. Right: Taxonomic composition of mapped contigs.

### Genetic map diagnostics

#### 1. IBD module

Before linkage mapping, IBD module (Rastas, 2017) was used to verify the relatedness between all individuals within families. In order to reduce computational needs, a subset of 10% of genetic markers was used. Heatmaps with IBD value plotted against each individual revealed that all samples were related, except from grandparents, which originated from different inbred lines (left bottom part of the plot).

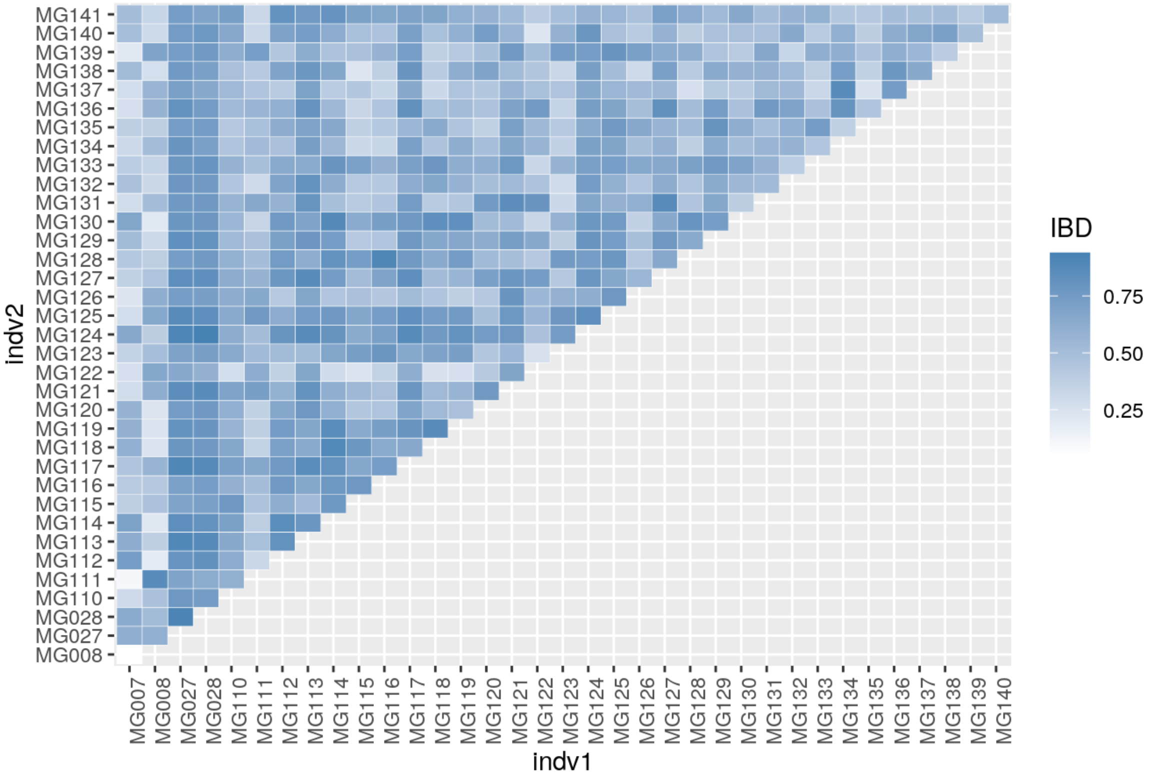


**Figure S2.** IBD calculated between individuals from family A.

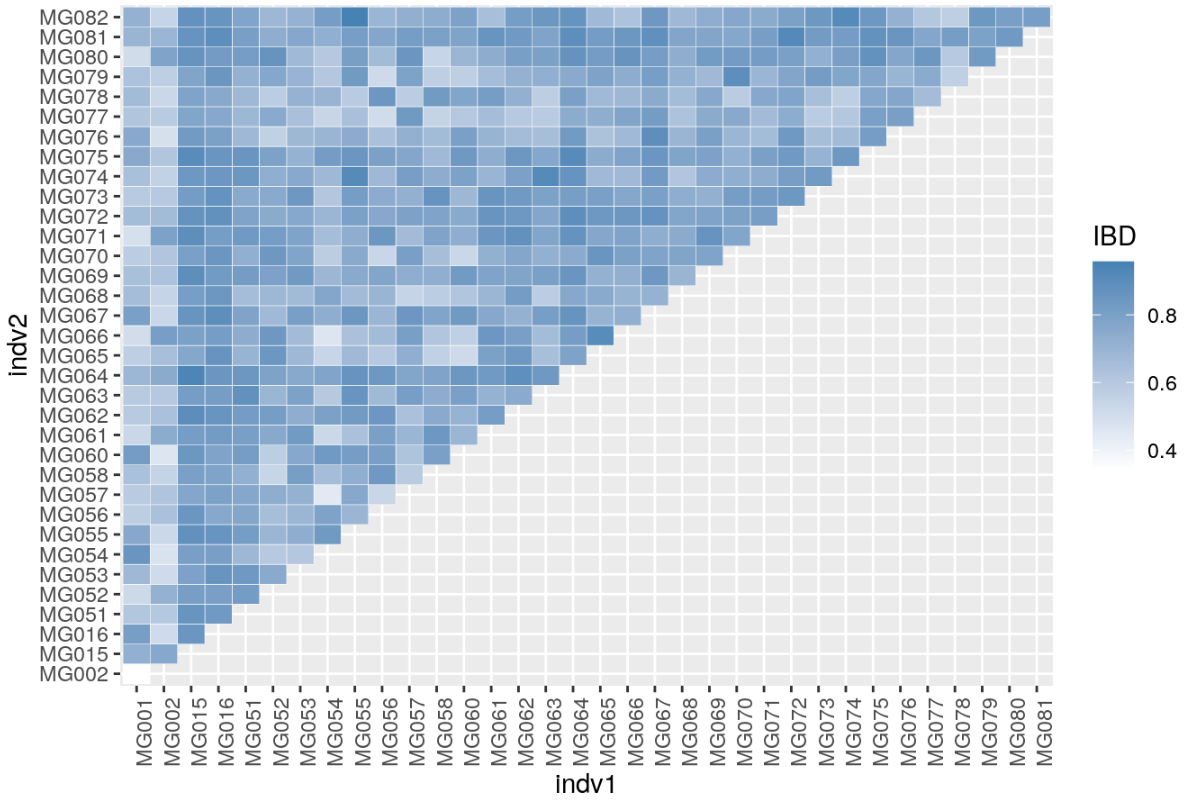


**Figure S3.** IBD calculated between individuals from family B.

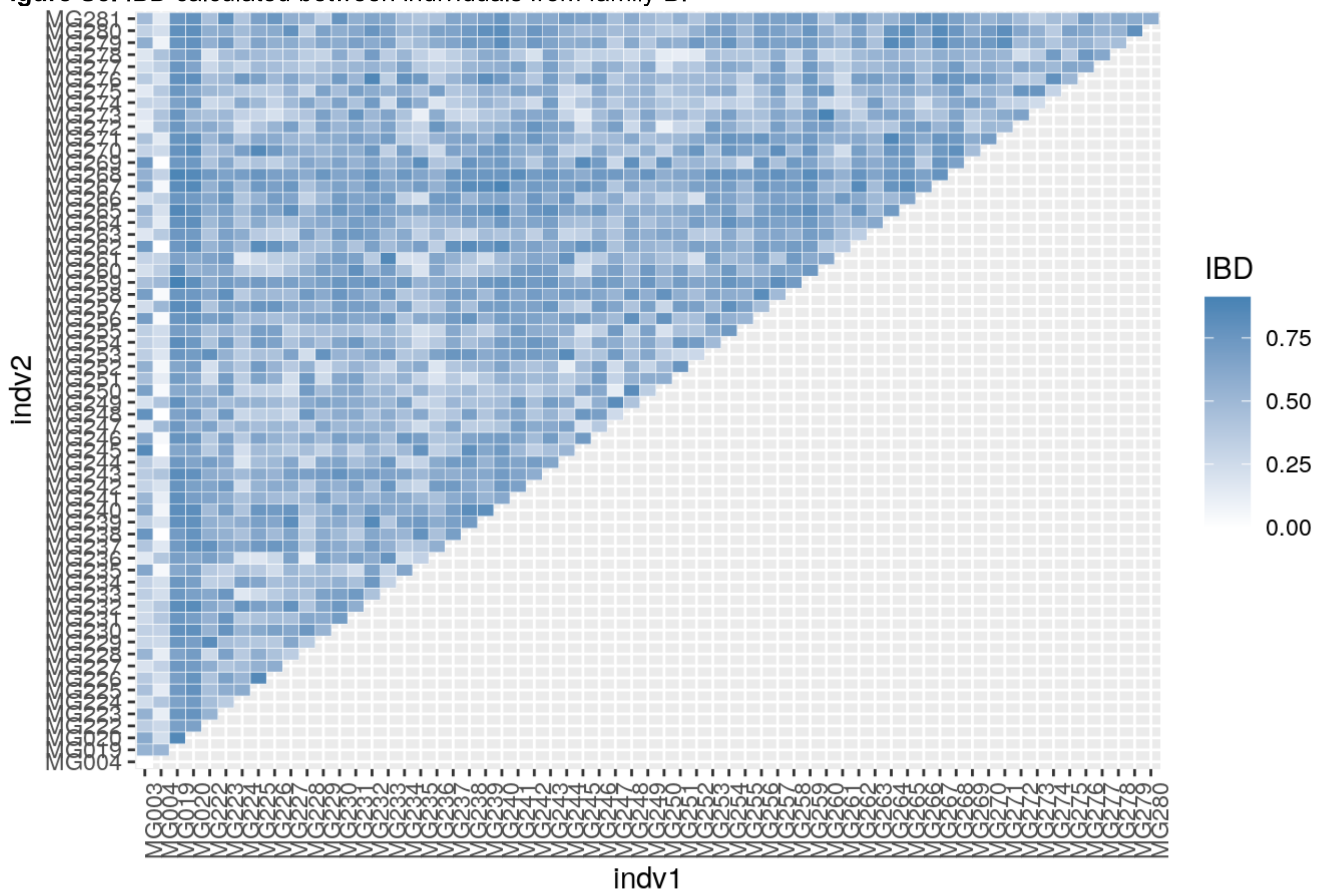


**Figure S4**. IBD calculated between individuals from family C.

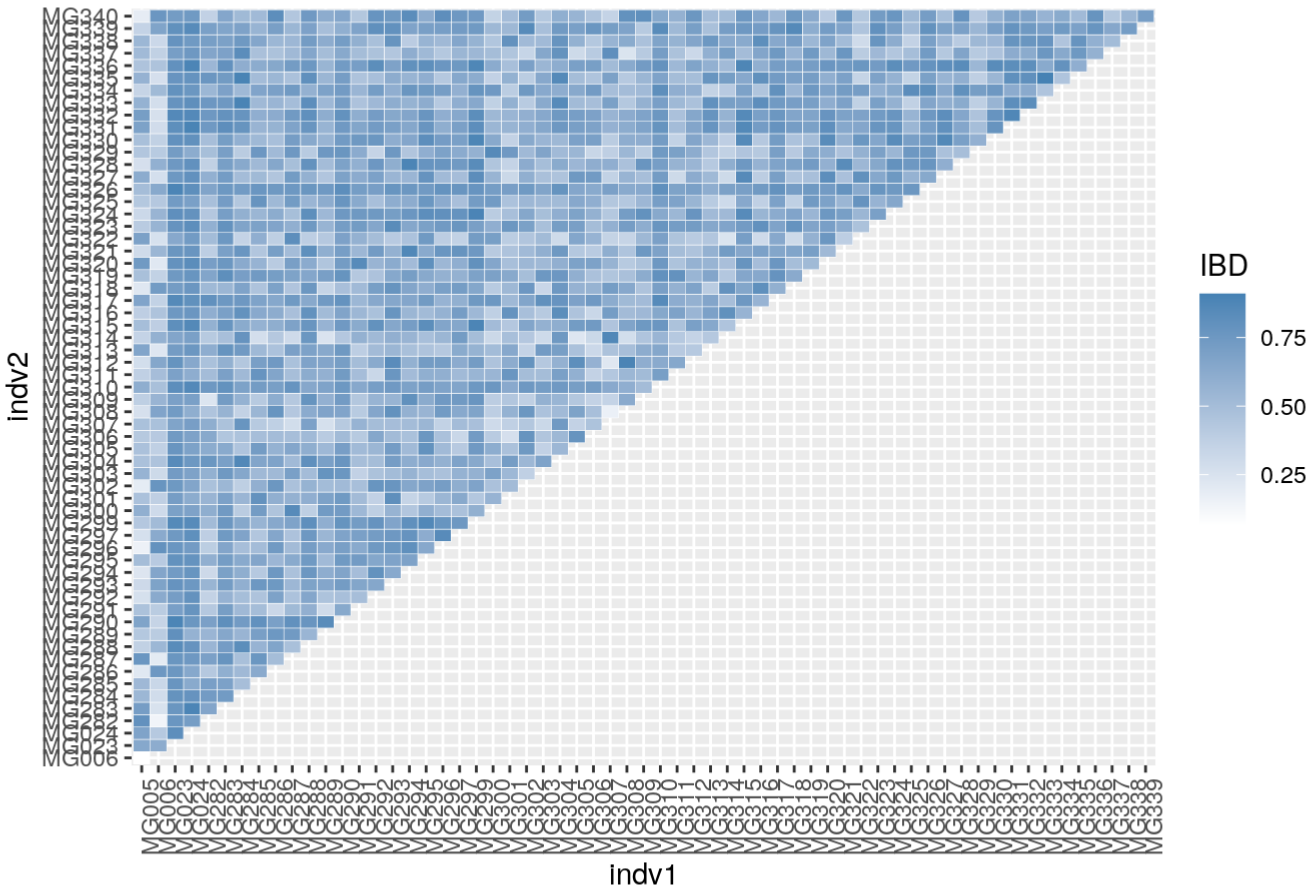


**Figure S5**. IBD calculated between individuals from family D.

### 2. Checking genotype missingness

To inspect the genotype missingness per sample, we called genotypes from 10 contigs, including 8 longest contigs from reference genome, and 2 sex-linked contigs using ANGSD (v. 0.937, Korneliussen et al., 2014) with parameters: -doPlink 2 -doGeno -4 -doPost 1 -doMajorMinor 1 -GL 1 -doCounts 1 -doMaf 2 -postCutoff 0.95 -SNP_pval 1e-6 -geno_minDepth 2. Next, the genotype file was used to check fraction of missing genotypes with plink (v. 1.9, Purcell et al., 2007) with –missing flag. The analysis revealed one outlier sample with 47% of missing genotypes, which was excluded from the subsequent analysis.

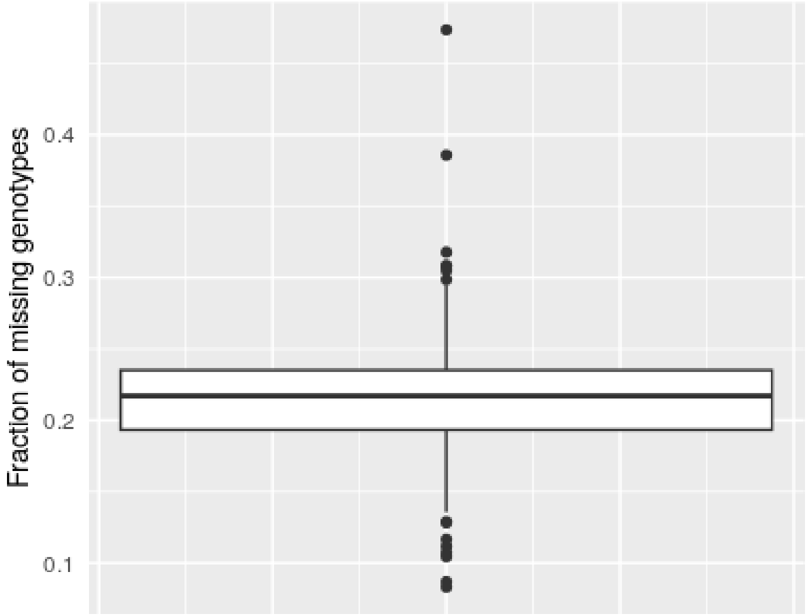


**Figure S6.** Boxplot displaying fraction of missing genotypes calculated for each sample. Central line is a median; box represent 25th and 75th quartiles; whiskers denote interquartile range.

### 3. Genetic map diagnostics with LMPlots

Map diagnostics was performed using LMPlot module. In LMPlots nodes represent distinct genetic positions with their size proportional to number of comprised genetic markers. Each line between nodes indicates one individual recombining between genetic positions.

LMPlots of sex-averaged maps used for genome anchoring were examined, and a single misjoin at linkage group 8 was observed (Supplementary Figure 6). To correct this error, the map was reordered wuth 60 iterations.

After anchoring the genome and genetic map evaluation, the maps were inspected again with LMPlots, and single misjoin was detected at chromosome 6 (Supplementary figure Y). The error was corrected again by manually equalizing the genetic distance value within the misjoin.

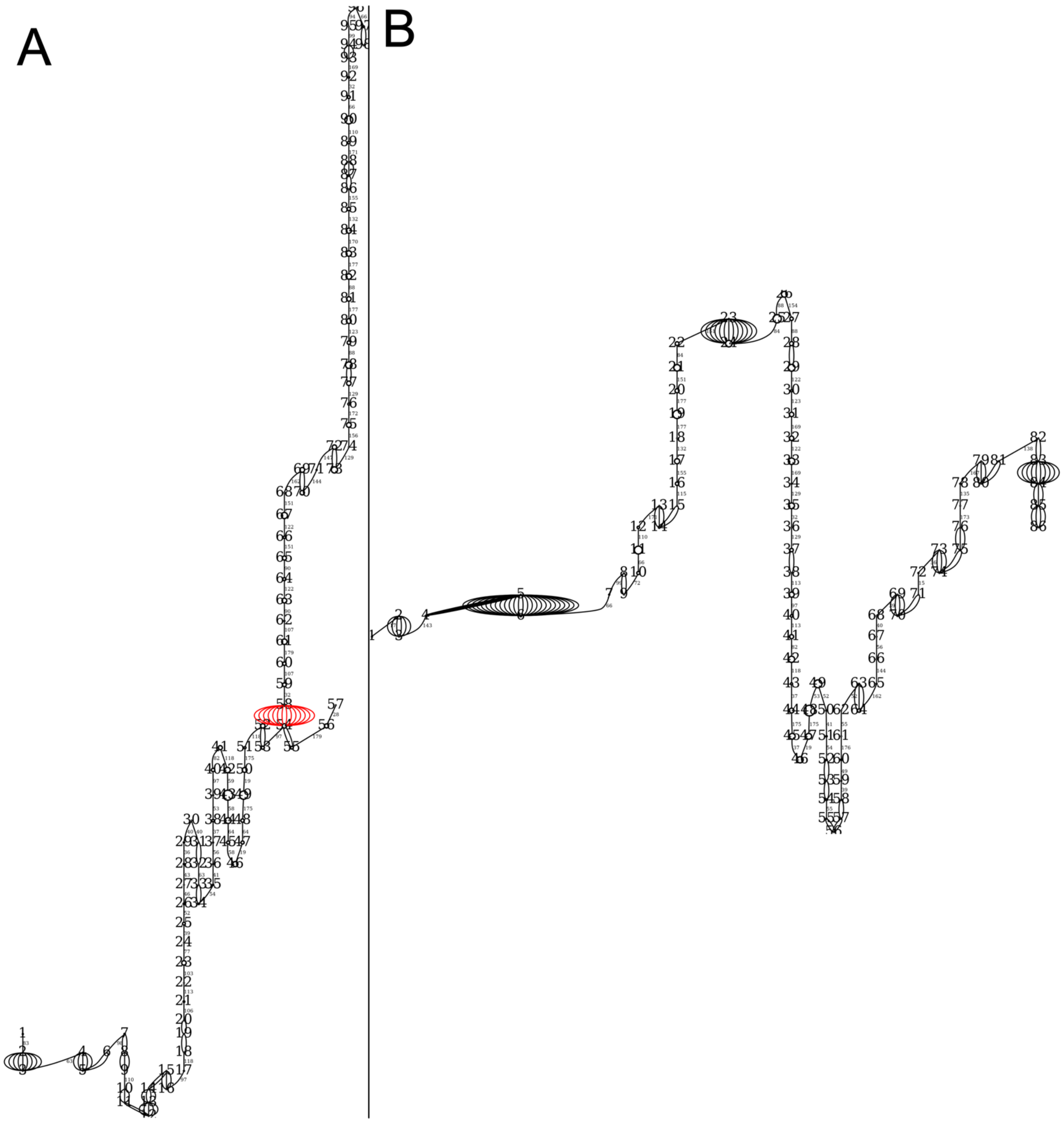


**Figure S7**. LMPlot of linkage group four before (A) and after (B) removing spurious recombinations (highlighted in red).

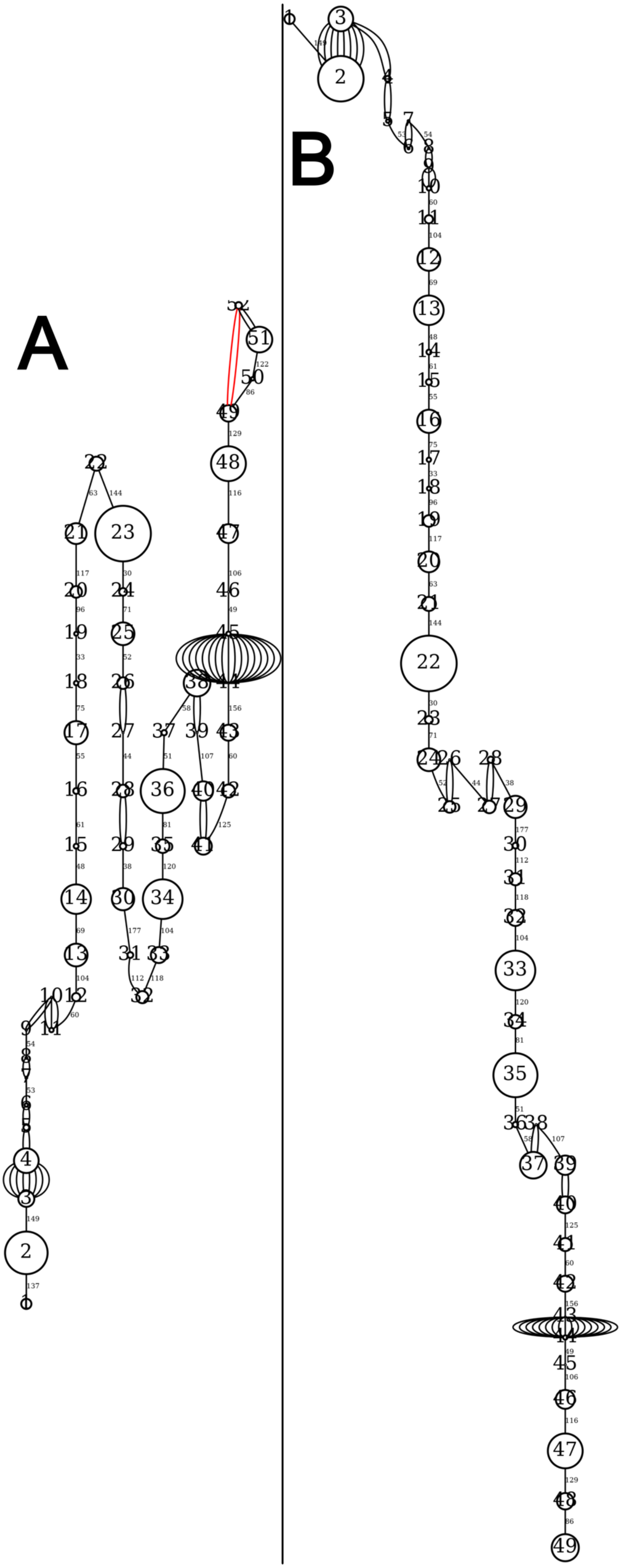


**Figure S8**. LMPlot of linkage group six before (A) and after (B) removing spurious recombinations (female map).

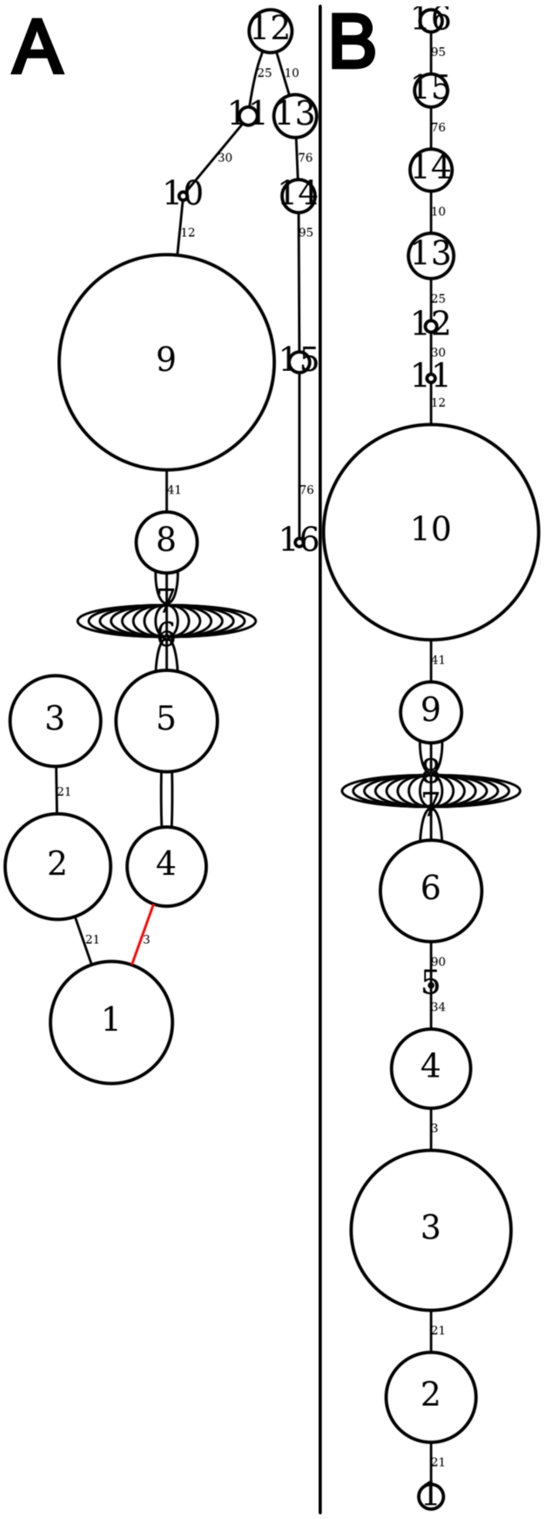


**Figure S9**. LMPlot of linkage group 5 before (A) and after (B) removing spurious recombinations (male map).

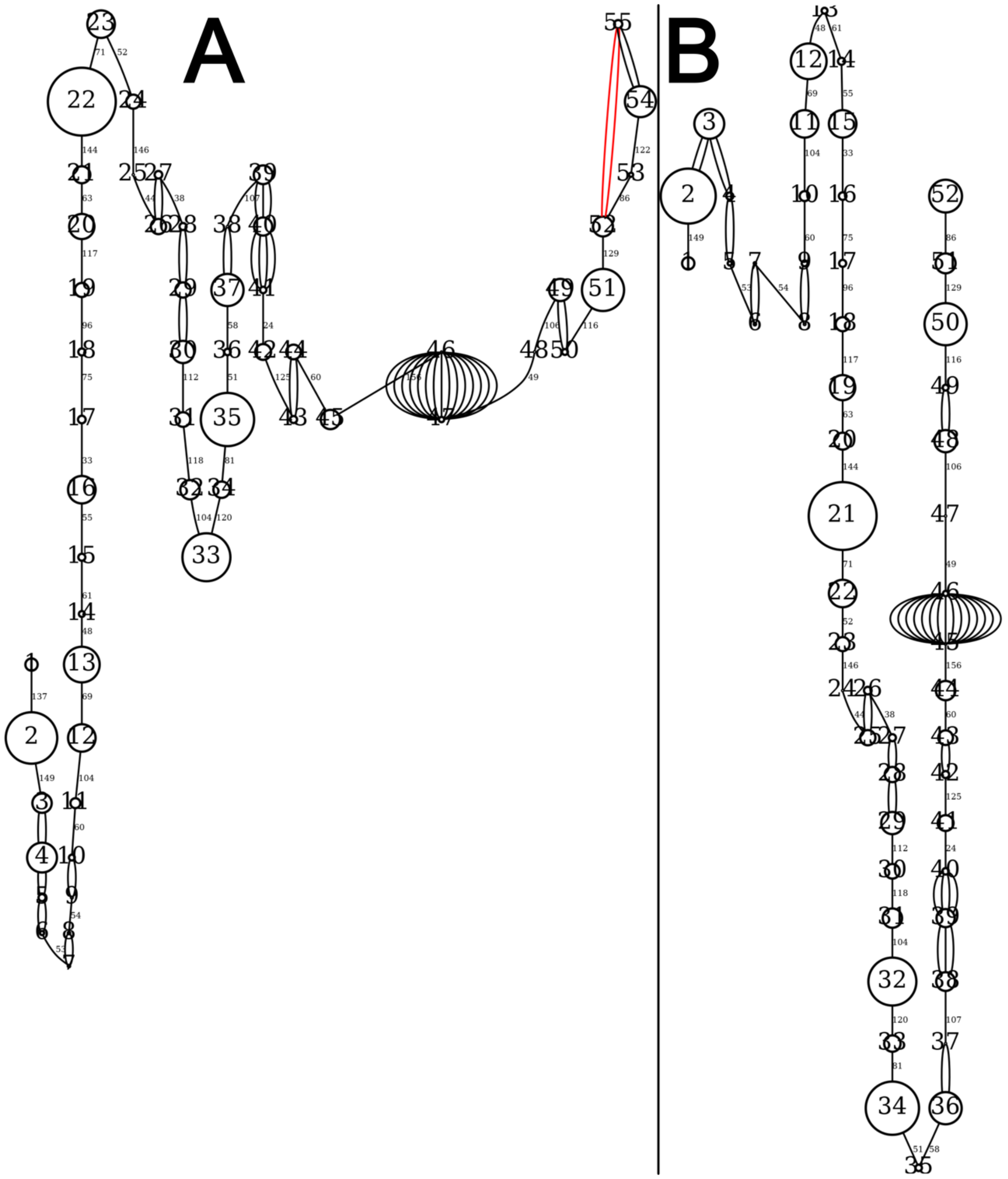


**Figure S10**. LMPlot of linkage group 6 before (A) and after (B) removing spurious recombinations (male map).

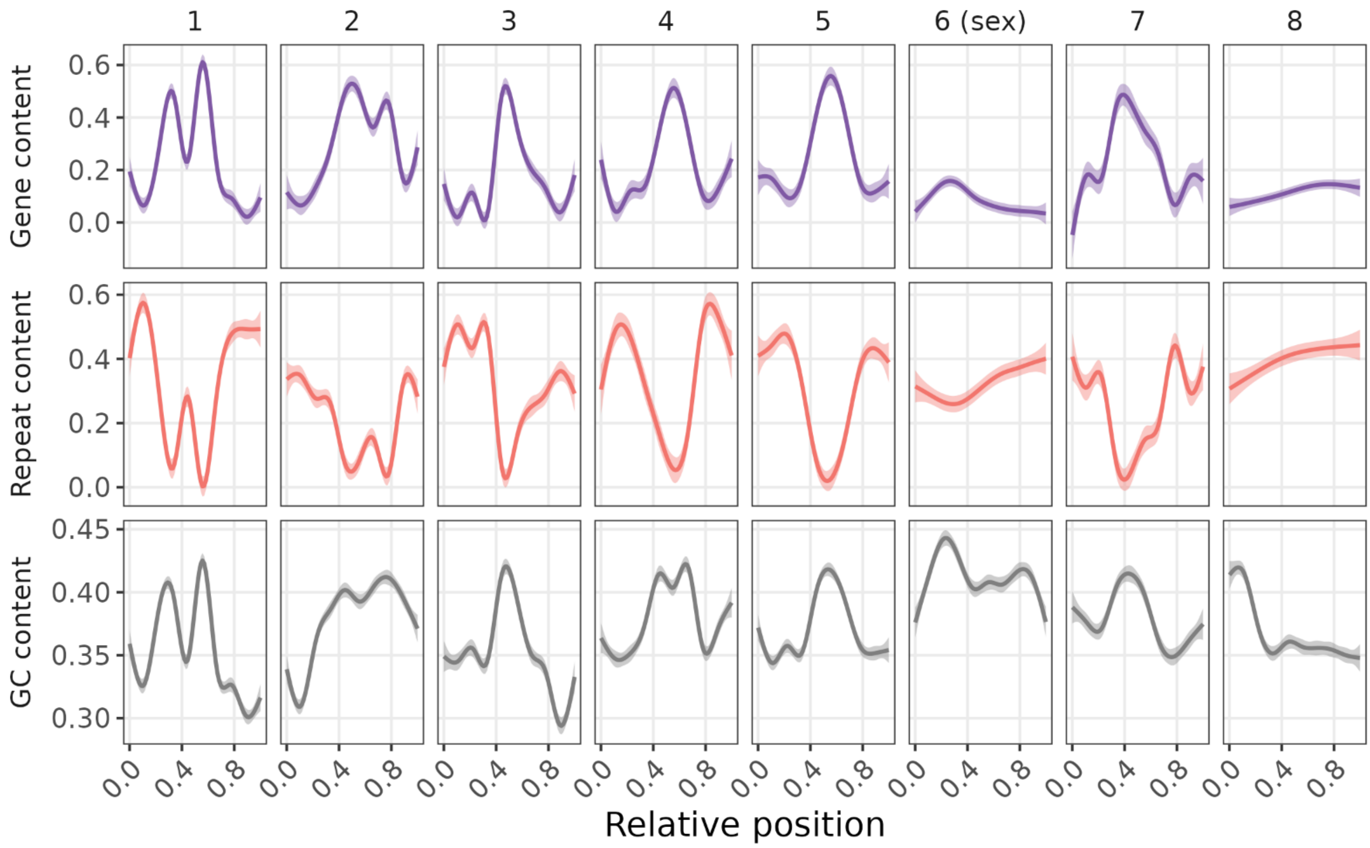


**Figure S11**. distribution of genomic features along bulb mite chromosomes measured in 50 kb windows. Upper panels: gene content, middle panels: repeat content and lower panels: GC nucleotides content.

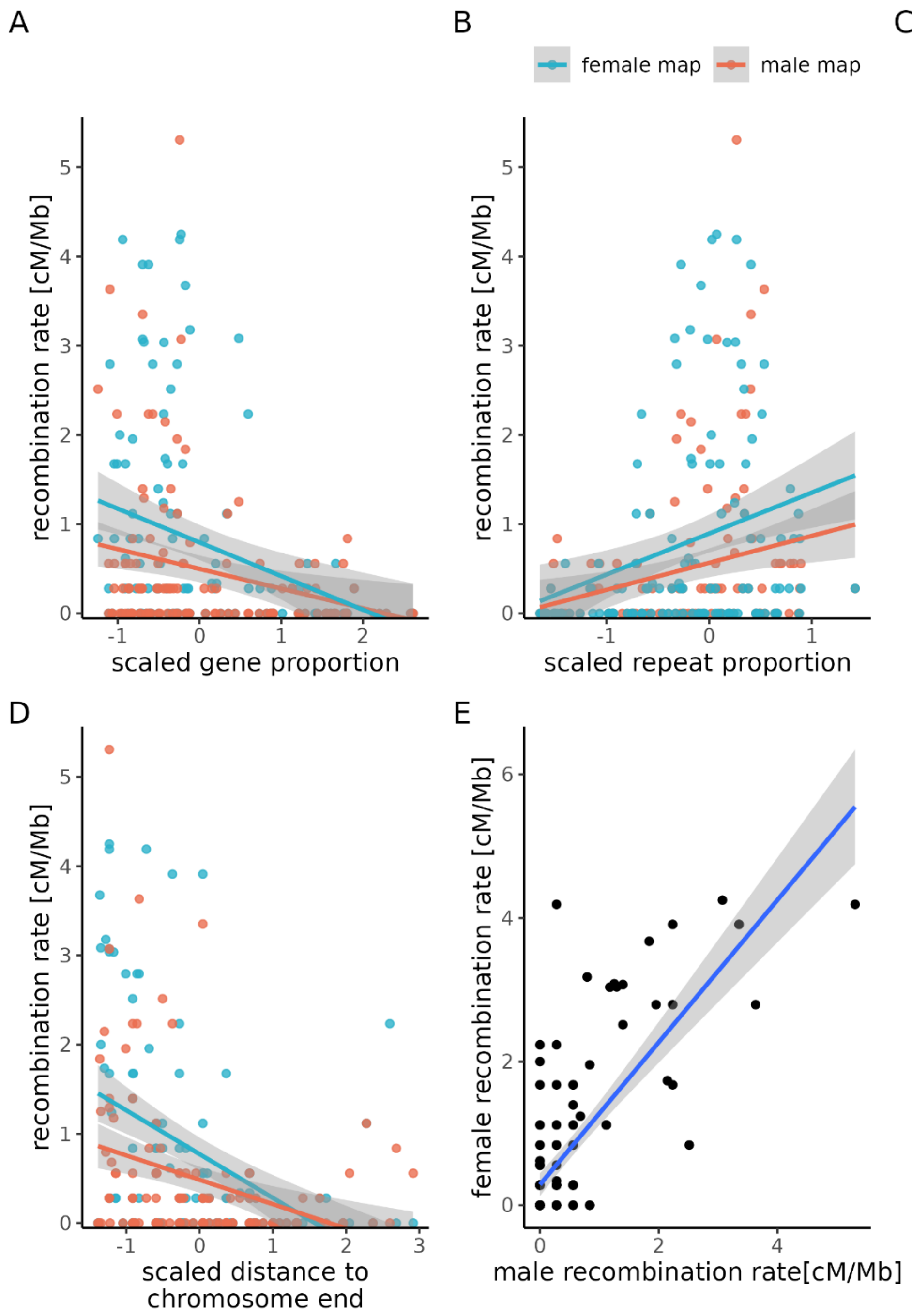


**Figure S12. Genomic features correlated with recombination rate in male and female linkage maps.** Panels A-D show the relationships between local recombination rate (cM/Mb) and scaled genomic features, with female map rates in blue and male map rates in red (solid lines represent linear regressions with 95% confidence intervals shaded in gray). Panel E: correlation between male and female recombination rates.

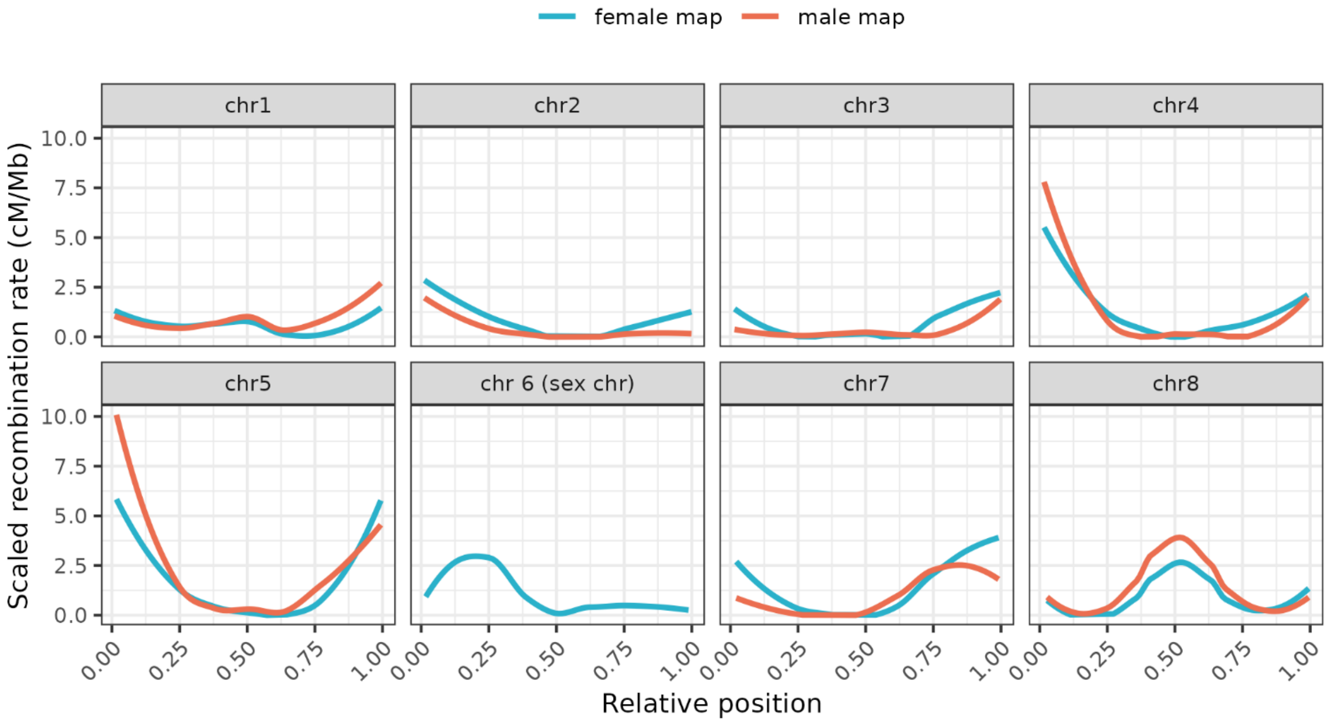


**Figure S13.** Sex-specific recombination rate for each chromosome calculated in 1Mb non-overlapping windows.

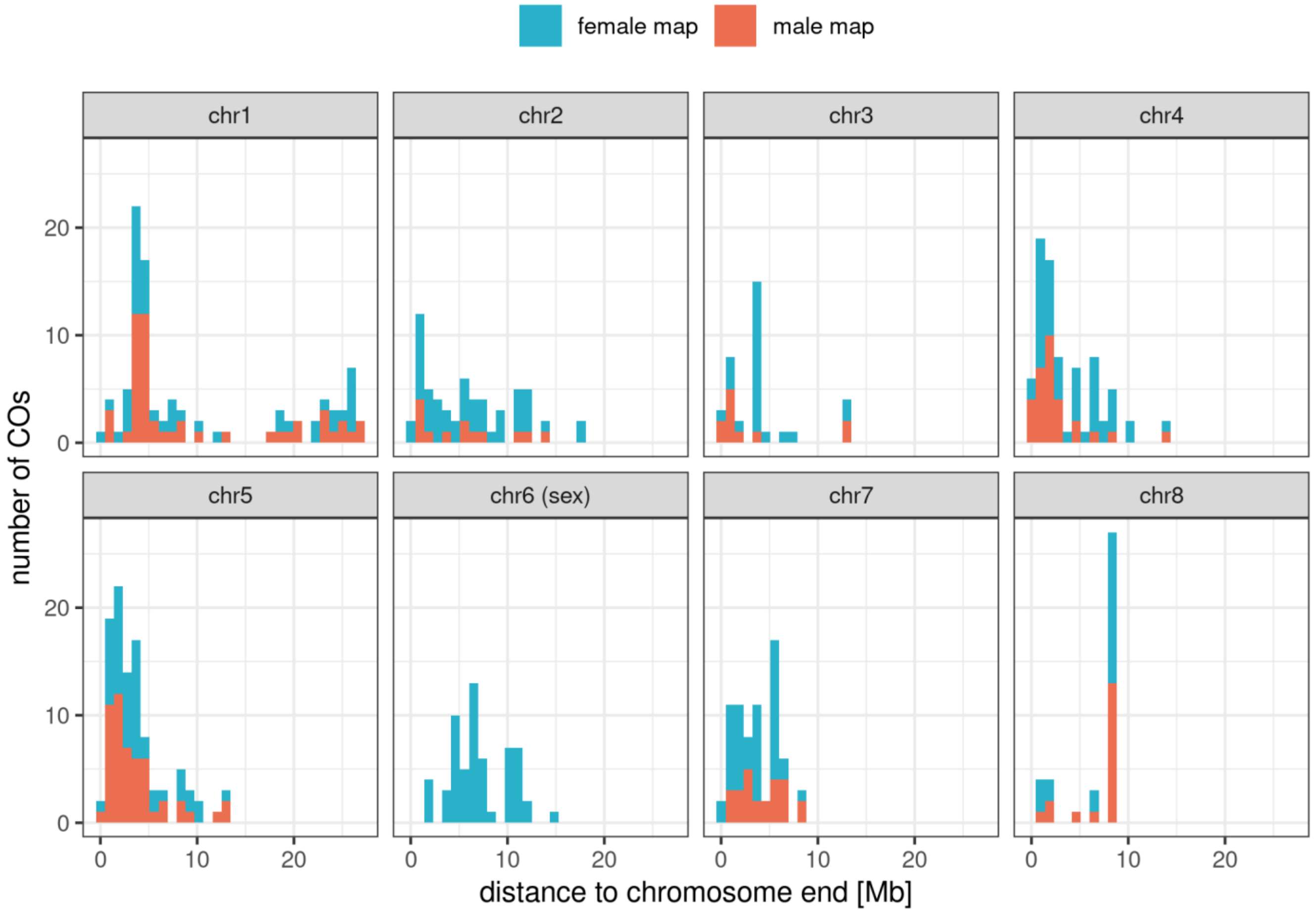


**Figure S14**. Histogram showing distance to chromosome ends for female (blue) and male (red) crossovers.

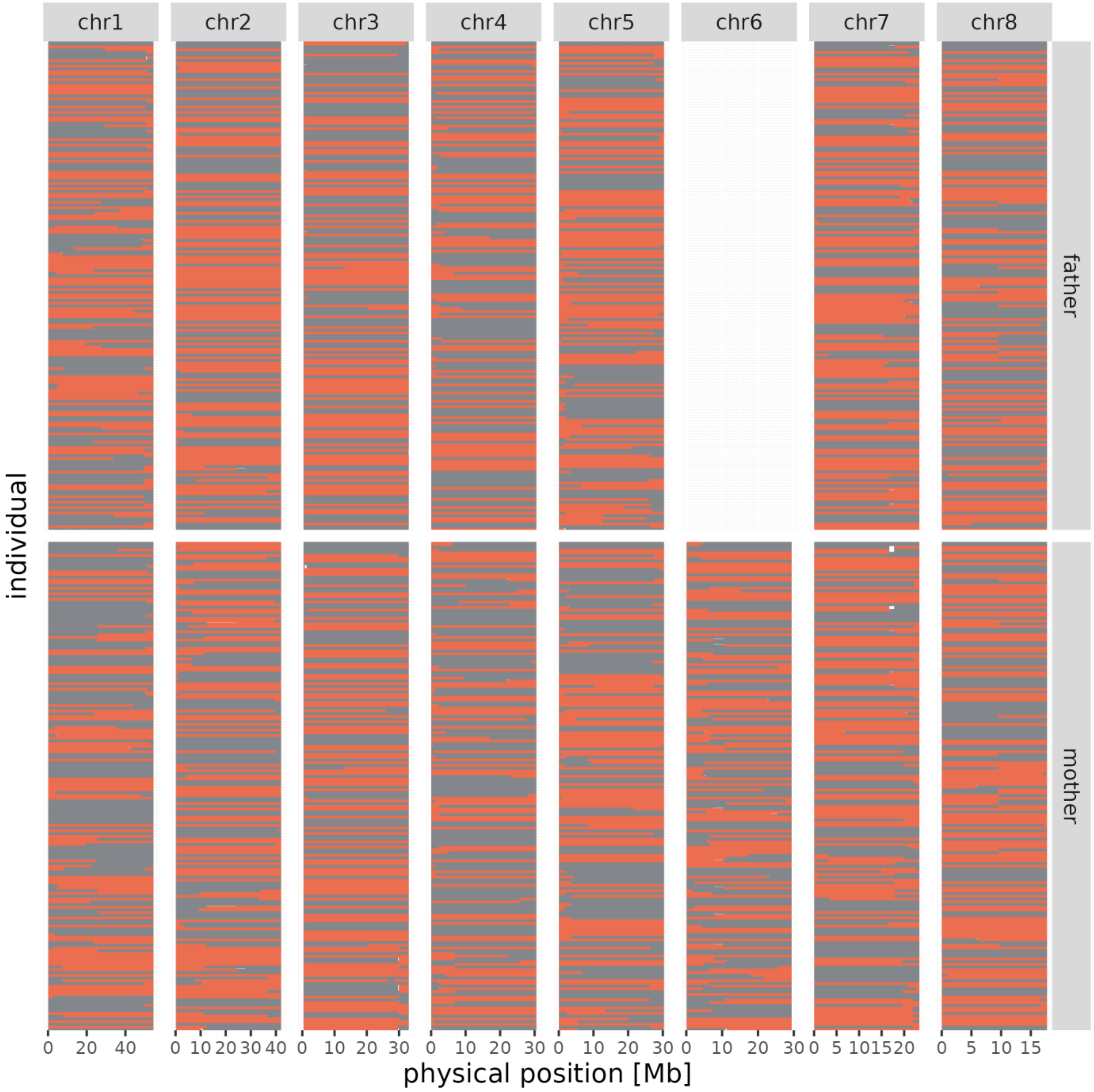


**Figure S15**. Chromosome phasing indicates the location of crossovers during paternal (top panel) and maternal (bottom panel) meiosis. Each row is one individual and X axis is chromosomal position in Mb. Small gaps in phasing are visualized by white horizontal bars.

**Table S1. Summary of sample sequencing metrics for each sample. The table presents the number of sequencing reads, number of mapped reads, mean coverage mean mapping quality and supplication rate.**

| sample name | number of reads | number of mapped reads | mean coverage | mean mapping quality | duplication rate | generation | family |
|---|---|---|---|---|---|---|---|
| MG001 | 78 473 482 | 60 985 514 | 26,4 | 44,8 | 28.8% | P | A |
| MG002 | 70 450 031 | 67 291 308 | 30,7 | 44,9 | 24.34% | P | A |
| MG015 | 68 378 771 | 59 095 033 | 26,6 | 44,9 | 22.64% | F1 | A |
| MG016 | 67 254 696 | 65 135 202 | 29,2 | 44,8 | 23.69% | F1 | A |
| MG051 | 23 077 774 | 22 006 735 | 10,1 | 45,2 | 16.57% | F2 | A |
| MG052 | 18 730 126 | 17 881 936 | 8,3 | 45,1 | 14.97% | F2 | A |
| MG053 | 22 990 347 | 22 274 406 | 10,3 | 45,1 | 16.55% | F2 | A |

| | | | | | | | |
|---|---|---|---|---|---|---|---|
| MG054 | 21 673 643 | 20 575 724 | 9,5 | 45,3 | 16.27% | F2 | A |
| MG055 | 52 172 879 | 49 855 863 | 23 | 45,2 | 21.5% | F2 | A |
| MG056 | 19 489 758 | 18 767 887 | 8,5 | 45,2 | 16.28% | F2 | A |
| MG057 | 26 076 818 | 25 056 882 | 11,6 | 45,4 | 16.86% | F2 | A |
| MG058 | 28 884 075 | 27 943 891 | 12,8 | 45,2 | 17.56% | F2 | A |
| MG060 | 18 641 883 | 17 675 048 | 8,1 | 45,3 | 15.17% | F2 | A |
| MG061 | 21 593 139 | 20 096 676 | 9,3 | 45,5 | 15.51% | F2 | A |
| MG062 | 27 868 049 | 25 644 940 | 11,7 | 45,1 | 17.23% | F2 | A |
| MG063 | 22 456 887 | 21 448 374 | 9,8 | 45,2 | 17.08% | F2 | A |
| MG064 | 39 324 306 | 38 171 910 | 17,3 | 44,9 | 19.12% | F2 | A |
| MG065 | 27 531 502 | 26 624 082 | 12 | 45,2 | 17.16% | F2 | A |
| MG066 | 31 625 580 | 30 240 392 | 13,8 | 45,2 | 17.83% | F2 | A |
| MG067 | 26 037 555 | 24 674 950 | 11,4 | 45,2 | 15.82% | F2 | A |
| MG068 | 25 173 891 | 24 046 678 | 11,1 | 45 | 17.03% | F2 | A |
| MG069 | 25 927 867 | 24 688 002 | 11,4 | 45 | 16.24% | F2 | A |
| MG070 | 20 966 446 | 20 006 467 | 9,3 | 45,2 | 14.65% | F2 | A |
| MG071 | 17 572 209 | 16 656 572 | 7,6 | 45 | 15% | F2 | A |
| MG072 | 24 535 423 | 23 683 527 | 11 | 45,1 | 16.55% | F2 | A |
| MG073 | 38 939 301 | 37 529 472 | 17,2 | 44,8 | 18.54% | F2 | A |
| MG074 | 29 953 723 | 28 481 205 | 13 | 44,9 | 17.08% | F2 | A |
| MG075 | 23 298 114 | 22 283 064 | 10,1 | 45 | 15.68% | F2 | A |
| MG076 | 19 329 872 | 18 046 027 | 8,4 | 45,4 | 14.56% | F2 | A |
| MG077 | 26 496 431 | 25 580 916 | 11,9 | 45,2 | 17.81% | F2 | A |
| MG078 | 20 212 760 | 19 461 511 | 9 | 45,2 | 15.27% | F2 | A |
| MG079 | 18 926 092 | 17 928 871 | 8,3 | 45,1 | 14.33% | F2 | A |
| MG080 | 24 528 315 | 23 351 203 | 10,9 | 45,2 | 15.84% | F2 | A |
| MG081 | 23 979 177 | 23 162 691 | 10,5 | 45 | 16.69% | F2 | A |
| MG082 | 32 693 184 | 30 844 253 | 14,3 | 45 | 17.15% | F2 | A |
| MG007 | 58 219 507 | 56 178 584 | 24,9 | 44,9 | 22.88% | P | B |
| MG008 | 54 249 331 | 48 854 772 | 20,5 | 44,7 | 24.06% | P | B |
| MG027 | 57 846 743 | 49 237 349 | 21,3 | 45,4 | 22.08% | F1 | B |
| MG028 | 60 366 241 | 54 718 719 | 24,5 | 45,7 | 22.22% | F1 | B |
| MG110 | 21 407 184 | 20 525 589 | 9,5 | 45,2 | 14.5% | F2 | B |
| MG111 | 23 726 266 | 22 833 582 | 10,6 | 45,7 | 15.99% | F2 | B |
| MG112 | 15 074 075 | 14 529 140 | 6,7 | 45,8 | 14.18% | F2 | B |
| MG113 | 23 125 900 | 22 121 078 | 10,3 | 45,9 | 16.2% | F2 | B |
| MG114 | 22 500 323 | 21 628 692 | 10,1 | 45,7 | 15.53% | F2 | B |
| MG115 | 18 404 986 | 17 116 919 | 7,9 | 45,2 | 14.65% | F2 | B |
| MG116 | 18 212 381 | 17 244 573 | 8 | 45,7 | 14.7% | F2 | B |
| MG117 | 40 104 492 | 38 171 166 | 17,8 | 46,1 | 18.05% | F2 | B |
| MG118 | 28 444 008 | 27 339 727 | 12,6 | 45,4 | 16.78% | F2 | B |
| MG119 | 27 603 636 | 26 637 316 | 12,4 | 45,8 | 16.49% | F2 | B |
| MG120 | 27 727 950 | 26 385 835 | 12,2 | 45,5 | 16.5% | F2 | B |
| MG121 | 22 739 158 | 21 935 941 | 10,3 | 45,7 | 15.06% | F2 | B |
| MG122 | 24 965 089 | 23 344 886 | 10,9 | 46,1 | 15.69% | F2 | B |
| MG123 | 19 794 126 | 18 881 796 | 8,7 | 45,3 | 15.77% | F2 | B |

| | | | | | | | |
|---|---|---|---|---|---|---|---|
| MG124 | 45 288 130 | 42 939 695 | 19,9 | 45,8 | 19.22% | F2 | B |
| MG125 | 40 920 800 | 38 307 423 | 17,7 | 45,9 | 18.57% | F2 | B |
| MG126 | 49 751 289 | 47 289 906 | 22 | 45,5 | 19.65% | F2 | B |
| MG127 | 49 383 457 | 47 200 170 | 22 | 46,1 | 19.7% | F2 | B |
| MG128 | 28 187 727 | 27 086 184 | 12,5 | 45,9 | 16.3% | F2 | B |
| MG129 | 16 610 062 | 15 988 319 | 7,4 | 46 | 14.25% | F2 | B |
| MG130 | 28 271 050 | 26 709 829 | 12,3 | 45,6 | 16.65% | F2 | B |
| MG131 | 19 603 668 | 18 747 902 | 8,8 | 46 | 14.17% | F2 | B |
| MG132 | 24 059 229 | 23 254 656 | 10,8 | 45,4 | 15.85% | F2 | B |
| MG133 | 41 308 700 | 39 507 993 | 18,4 | 45,6 | 19.75% | F2 | B |
| MG134 | 35 888 679 | 33 933 433 | 15,8 | 45,9 | 17.85% | F2 | B |
| MG135 | 26 963 311 | 26 094 115 | 12,1 | 45,4 | 17.28% | F2 | B |
| MG136 | 28 631 856 | 27 774 051 | 12,7 | 45,9 | 17.91% | F2 | B |
| MG137 | 27 965 714 | 26 330 660 | 12,2 | 45,6 | 16.92% | F2 | B |
| MG138 | 21 956 993 | 21 146 517 | 9,8 | 45,7 | 15.82% | F2 | B |
| MG139 | 20 689 248 | 19 607 748 | 9,2 | 45,9 | 14.95% | F2 | B |
| MG140 | 26 701 211 | 25 494 419 | 11,9 | 45,1 | 16.24% | F2 | B |
| MG141 | 47 847 927 | 46 024 953 | 21,3 | 45,6 | 20.48% | F2 | B |
| MG003 | 50 170 764 | 48 242 344 | 20,9 | 44,4 | 23.42% | P | C |
| MG004 | 59 751 855 | 58 089 236 | 26 | 44,7 | 24% | P | C |
| MG019 | 65 367 023 | 62 062 017 | 28 | 44,9 | 23.42% | F1 | C |
| MG020 | 65 137 900 | 62 227 600 | 28,2 | 45,1 | 23.24% | F1 | C |
| MG222 | 20 953 848 | 19 507 180 | 8,8 | 45,1 | 25.87% | F2 | C |
| MG223 | 23 578 614 | 22 406 403 | 10,2 | 44,7 | 29.34% | F2 | C |
| MG224 | 26 090 721 | 25 061 275 | 11,5 | 45,1 | 26.79% | F2 | C |
| MG225 | 24 967 955 | 23 462 043 | 10,8 | 45,1 | 29.53% | F2 | C |
| MG226 | 23 172 363 | 22 248 602 | 10,2 | 45,2 | 26.3% | F2 | C |
| MG227 | 20 917 783 | 20 071 886 | 9,3 | 45,1 | 27.44% | F2 | C |
| MG228 | 30 347 011 | 28 687 617 | 12,9 | 44,6 | 29.77% | F2 | C |
| MG229 | 22 712 970 | 21 584 856 | 9,8 | 44,8 | 28.45% | F2 | C |
| MG230 | 22 982 926 | 21 879 400 | 10 | 45,1 | 26.71% | F2 | C |
| MG231 | 11 515 213 | 9 189 300 | 4,2 | 44,5 | 27.46% | F2 | C |
| MG232 | 25 807 206 | 24 717 554 | 11,4 | 45,3 | 26.94% | F2 | C |
| MG233 | 27 962 546 | 26 548 362 | 12,2 | 44,6 | 27.57% | F2 | C |
| MG234 | 17 195 720 | 15 519 332 | 7,2 | 44,9 | 24.35% | F2 | C |
| MG235 | 30 339 650 | 27 563 607 | 12,6 | 44,7 | 27.89% | F2 | C |
| MG236 | 27 822 146 | 26 631 611 | 12,2 | 44,5 | 27.76% | F2 | C |
| MG237 | 23 415 542 | 21 707 591 | 10 | 44,9 | 28.11% | F2 | C |
| MG238 | 27 411 535 | 26 199 132 | 12 | 44,7 | 26.56% | F2 | C |
| MG239 | 26 935 027 | 25 637 718 | 11,8 | 44,9 | 27.77% | F2 | C |
| MG240 | 23 249 518 | 21 969 494 | 10,1 | 45 | 24.88% | F2 | C |
| MG241 | 22 069 797 | 21 062 177 | 9,7 | 44,5 | 26.52% | F2 | C |
| MG242 | 30 176 871 | 28 631 508 | 13,2 | 44,4 | 26.25% | F2 | C |
| MG243 | 21 186 383 | 20 256 988 | 9,3 | 44,8 | 26.58% | F2 | C |
| MG244 | 31 615 644 | 29 980 484 | 13,8 | 44,8 | 28.05% | F2 | C |
| MG245 | 29 890 669 | 27 519 849 | 12,6 | 44,7 | 29.66% | F2 | C |

| | | | | | | | |
|---|---|---|---|---|---|---|---|
| MG246 | 23 231 281 | 21 798 354 | 10 | 44,7 | 25.43% | F2 | C |
| MG247 | 20 636 134 | 19 683 217 | 9,1 | 44,8 | 26.64% | F2 | C |
| MG248 | 22 148 235 | 21 116 763 | 9,7 | 44,4 | 25.61% | F2 | C |
| MG249 | 24 293 815 | 23 229 690 | 10,7 | 44,8 | 26.94% | F2 | C |
| MG250 | 18 810 856 | 17 819 515 | 8,1 | 44,4 | 24.63% | F2 | C |
| MG251 | 18 951 668 | 17 879 449 | 8,2 | 44,4 | 25.64% | F2 | C |
| MG252 | 24 447 656 | 22 814 256 | 10,5 | 44,3 | 25.39% | F2 | C |
| MG253 | 22 179 515 | 20 951 949 | 9,6 | 45,1 | 26.43% | F2 | C |
| MG254 | 30 750 904 | 29 112 706 | 13,4 | 44,3 | 29.22% | F2 | C |
| MG255 | 26 362 657 | 25 233 520 | 11,7 | 44,6 | 30.6% | F2 | C |
| MG256 | 28 420 468 | 27 029 879 | 12,4 | 44,8 | 26.24% | F2 | C |
| MG257 | 20 788 638 | 19 829 477 | 9,2 | 44,5 | 25.98% | F2 | C |
| MG258 | 24 976 601 | 23 888 865 | 10,9 | 44,5 | 25.76% | F2 | C |
| MG259 | 20 613 282 | 19 715 002 | 9,1 | 44,8 | 26.1% | F2 | C |
| MG260 | 27 059 661 | 25 755 644 | 11,8 | 44,2 | 27.07% | F2 | C |
| MG261 | 21 367 913 | 20 387 357 | 9,3 | 44,4 | 28.25% | F2 | C |
| MG262 | 26 738 985 | 25 526 041 | 11,7 | 44,8 | 25.99% | F2 | C |
| MG263 | 24 944 566 | 23 871 172 | 10,9 | 44,8 | 27.68% | F2 | C |
| MG264 | 27 836 336 | 26 159 287 | 12 | 44,5 | 25.87% | F2 | C |
| MG265 | 23 872 840 | 22 880 051 | 10,5 | 44,8 | 27.16% | F2 | C |
| MG266 | 19 541 022 | 18 687 145 | 8,5 | 44,8 | 25.44% | F2 | C |
| MG267 | 28 921 734 | 27 544 315 | 12,6 | 44,9 | 28.09% | F2 | C |
| MG268 | 23 216 240 | 22 076 893 | 10,1 | 44,6 | 25.69% | F2 | C |
| MG269 | 26 643 149 | 25 152 955 | 11,5 | 44,7 | 28.45% | F2 | C |
| MG270 | 23 810 209 | 22 709 151 | 10,5 | 44,9 | 25.63% | F2 | C |
| MG271 | 22 708 780 | 21 535 021 | 9,9 | 44,8 | 27.37% | F2 | C |
| MG272 | 27 899 243 | 26 623 061 | 12,3 | 44,6 | 27.95% | F2 | C |
| MG273 | 30 266 019 | 28 973 605 | 13,3 | 44,4 | 30.09% | F2 | C |
| MG274 | 28 520 076 | 26 650 678 | 12,2 | 44,7 | 26.63% | F2 | C |
| MG275 | 28 067 835 | 27 021 108 | 12,4 | 44,5 | 28.67% | F2 | C |
| MG276 | 24 003 734 | 23 022 979 | 10,5 | 44,7 | 25.72% | F2 | C |
| MG277 | 21 792 187 | 21 003 148 | 9,6 | 44,6 | 27.13% | F2 | C |
| MG278 | 24 644 411 | 23 647 350 | 10,9 | 44,9 | 25.95% | F2 | C |
| MG279 | 28 683 411 | 27 443 695 | 12,6 | 44,8 | 28.31% | F2 | C |
| MG280 | 24 690 186 | 23 656 031 | 10,9 | 44,8 | 25.47% | F2 | C |
| MG281 | 23 547 032 | 22 574 782 | 10,3 | 44,9 | 27.12% | F2 | C |
| MG005 | 58 480 478 | 55 460 759 | 25,2 | 44,5 | 22.32% | P | D |
| MG006 | 57 899 290 | 54 387 857 | 24,6 | 45,1 | 22.73% | P | D |
| MG023 | 67 437 922 | 61 336 956 | 28,1 | 45,3 | 23.19% | F1 | D |
| MG024 | 68 944 076 | 65 266 260 | 30,1 | 45,2 | 23.98% | F1 | D |
| MG282 | 19 888 747 | 18 882 199 | 8,7 | 44,7 | 25.16% | F2 | D |
| MG283 | 21 345 036 | 20 420 912 | 9,4 | 44,4 | 27.1% | F2 | D |
| MG284 | 21 313 443 | 18 282 303 | 8,4 | 44,5 | 25.43% | F2 | D |
| MG285 | 25 190 448 | 24 051 248 | 11,1 | 44,4 | 27.88% | F2 | D |
| MG286 | 22 060 716 | 21 146 199 | 9,7 | 44,6 | 25.76% | F2 | D |
| MG287 | 28 294 353 | 26 996 885 | 12,4 | 44,5 | 27.98% | F2 | D |

| | | | | | | | |
|---|---|---|---|---|---|---|---|
| MG288 | 22 400 058 | 21 530 422 | 9,9 | 44,7 | 25.25% | F2 | D |
| MG289 | 22 973 855 | 21 739 696 | 10 | 44,4 | 26.75% | F2 | D |
| MG290 | 23 997 990 | 22 655 946 | 10,4 | 44,5 | 25.6% | F2 | D |
| MG291 | 24 654 268 | 23 521 784 | 10,9 | 44,5 | 27.14% | F2 | D |
| MG292 | 24 221 831 | 21 327 664 | 9,8 | 44,8 | 26.42% | F2 | D |
| MG293 | 24 166 512 | 22 864 907 | 10,6 | 44,7 | 27.94% | F2 | D |
| MG294 | 21 302 659 | 20 462 935 | 9,5 | 44,6 | 25.73% | F2 | D |
| MG296 | 22 848 460 | 21 705 583 | 10 | 44,6 | 25.36% | F2 | D |
| MG297 | 25 290 465 | 24 042 840 | 11,1 | 44,6 | 27.74% | F2 | D |
| MG299 | 28 335 182 | 27 256 438 | 12,5 | 44,8 | 27.61% | F2 | D |
| MG300 | 20 765 394 | 19 750 828 | 9 | 44,1 | 25.61% | F2 | D |
| MG301 | 27 984 563 | 25 536 578 | 11,7 | 44,5 | 27.54% | F2 | D |
| MG302 | 21 836 598 | 20 971 240 | 9,7 | 45,1 | 25.15% | F2 | D |
| MG303 | 28 287 284 | 26 790 697 | 12,3 | 44,2 | 27.67% | F2 | D |
| MG304 | 19 361 046 | 18 414 766 | 8,5 | 44,5 | 25.1% | F2 | D |
| MG305 | 25 985 275 | 24 961 586 | 11,5 | 45 | 28.25% | F2 | D |
| MG306 | 24 124 651 | 22 735 121 | 10,5 | 44,9 | 25.53% | F2 | D |
| MG307 | 39 302 554 | 36 789 213 | 17 | 44,9 | 29.49% | F2 | D |
| MG308 | 22 475 012 | 21 459 343 | 9,9 | 44,7 | 24.71% | F2 | D |
| MG309 | 28 604 267 | 26 074 272 | 12,1 | 44,3 | 26.51% | F2 | D |
| MG310 | 22 079 449 | 20 756 224 | 9,6 | 44,8 | 24.52% | F2 | D |
| MG311 | 26 531 983 | 25 462 214 | 11,7 | 44,4 | 27.65% | F2 | D |
| MG312 | 23 048 850 | 22 246 988 | 10,3 | 44,9 | 28.01% | F2 | D |
| MG313 | 29 089 823 | 27 936 164 | 12,9 | 44,5 | 30.67% | F2 | D |
| MG314 | 20 894 151 | 19 986 263 | 9,3 | 44,7 | 25.8% | F2 | D |
| MG315 | 28 448 865 | 27 007 095 | 12,5 | 44,6 | 27.81% | F2 | D |
| MG316 | 26 700 958 | 25 411 386 | 11,7 | 44,8 | 27.46% | F2 | D |
| MG317 | 28 313 907 | 25 438 312 | 11,7 | 44,7 | 28.59% | F2 | D |
| MG318 | 29 261 933 | 27 324 895 | 12,7 | 44,9 | 27.13% | F2 | D |
| MG319 | 28 227 386 | 26 597 567 | 12,3 | 44,4 | 28.76% | F2 | D |
| MG320 | 24 812 432 | 23 671 088 | 10,9 | 44,6 | 26.29% | F2 | D |
| MG321 | 24 591 889 | 23 425 545 | 10,9 | 44,6 | 27.16% | F2 | D |
| MG322 | 24 594 174 | 23 304 777 | 10,8 | 44,3 | 26.35% | F2 | D |
| MG323 | 24 867 066 | 23 744 733 | 11 | 44,5 | 27.74% | F2 | D |
| MG324 | 25 712 211 | 24 484 251 | 11,4 | 44,8 | 25.82% | F2 | D |
| MG325 | 25 589 204 | 24 004 134 | 11,1 | 44,4 | 27.08% | F2 | D |
| MG326 | 21 079 447 | 20 160 387 | 9,3 | 44,6 | 24.79% | F2 | D |
| MG327 | 24 993 232 | 23 961 477 | 11,1 | 44,7 | 27.36% | F2 | D |
| MG328 | 22 812 320 | 21 857 132 | 10,1 | 44,5 | 26.22% | F2 | D |
| MG329 | 25 624 691 | 24 486 090 | 11,3 | 43,9 | 28.16% | F2 | D |
| MG330 | 19 356 761 | 17 959 967 | 8,3 | 44,6 | 24.21% | F2 | D |
| MG331 | 21 508 807 | 20 463 381 | 9,5 | 44,8 | 26.22% | F2 | D |
| MG332 | 23 988 377 | 22 924 185 | 10,6 | 44,6 | 28.43% | F2 | D |
| MG333 | 32 943 241 | 31 399 928 | 14,5 | 44,6 | 31.15% | F2 | D |
| MG334 | 23 053 142 | 22 023 182 | 10,2 | 44,7 | 24.92% | F2 | D |
| MG335 | 24 235 793 | 22 895 747 | 10,6 | 44,5 | 26.45% | F2 | D |

| | | | | | | | |
|---|---|---|---|---|---|---|---|
| MG336 | 22 526 702 | 21 366 658 | 9,9 | 44,6 | 25.52% | F2 | D |
| MG337 | 22 381 220 | 21 343 840 | 9,8 | 44,7 | 28.08% | F2 | D |
| MG338 | 15 023 630 | 14 217 179 | 6,6 | 44,8 | 23.26% | F2 | D |
| MG339 | 23 630 058 | 22 528 438 | 10,5 | 44,9 | 25.96% | F2 | D |
| MG340 | 22 057 788 | 20 963 276 | 9,7 | 44,6 | 24.81% | F2 | D |


Korneliussen, T. S., Albrechtsen, A., & Nielsen, R. (2014). Angsd. *BMC Bioinformatics*, *15*(1), 1–13.

Purcell, S., Neale, B., Todd-Brown, K., Thomas, L., Ferreira, M. A. R., Bender, D., Maller, J., Sklar, P., De Bakker, P. I. W., Daly, M. J., & Sham, P. C. (2007). PLINK: A tool set for whole-genome association and population-based linkage analyses. *American Journal of Human Genetics*, *81*(3), 559–575. https://doi.org/10.1086/519795

Rastas, P. (2017). Lep-MAP3: Robust linkage mapping even for low-coverage whole genome sequencing data. *Bioinformatics*, *33*(23), 3726–3732. https://doi.org/10.1093/bioinformatics/btx494
